# Dynamics of the distribution of fitness effects during adaptation

**DOI:** 10.1101/2024.10.09.617468

**Authors:** Tenoch Morales, Abigail Kushnir, Lindi M. Wahl

## Abstract

Mutations drive adaptive evolution due to their heritable effects on fitness. Empirical measures of the distribution of fitness effects of new mutations (the DFE) have been increasingly successful, and have recently highlighted the fact that the DFE changes during adaptation. Here, we analyze these dynamic changes to the DFE during a simplified adaptive process: an adaptive walk across additive and multiplicative fitness landscapes. First, we derive analytical approximations for the underlying fitness distributions of alleles present in the genotype and available through mutation and use these to derive expressions for the DFE at each step of the adaptive walk. We then confirm these predictions with independent simulations that relax several simplifying assumptions made in the analysis. As expected, our analysis predicts that as adaptation proceeds, the DFE is reshaped through an increase in deleterious mutations (a shift to the left). Surprisingly, this change occurs through different mechanisms depending on the number of alleles available per site: for a small number of available alleles, we observe a depletion of high-fitness alleles available through mutation as expected, however for a large number of alleles we observe that adaptation may be more limited by the availability of low-fitness alleles to be replaced, rather than by the availability of high-fitness alleles to replace them.

## 1 Introduction

Mutations are the building blocks of adaptive evolution, due to their heritable effects on organismal fitness. The fitness effects of mutations range from beneficial, leading to better adapted organisms, to mildly or strongly deleterious or even lethal (Eyre-Walker and Keightley, 2007). The distribution of fitness effects of new mutations (the DFE) is therefore a key factor in predicting the course of evolution (Wortel et al., 2023), since it quantifies the abundance of mutations with specific effect sizes.

Empirical measures of the DFE have been increasingly successful in the last 20 years, gaining resolution as new techniques in both DNA sequencing and fitness assays have allowed for fuller explorations of genotypic space. Examples include genome-wide DFEs measured in mutation accumulation experiments in yeast (Zeyl and DeVisser, 2001), bacteria (Kassen and Bataillon, 2006; Sane et al., 2023), viruses (Sanjuán et al., 2004), and uni- and multi-cellular plants (Böndel et al., 2019; Weng et al., 2021), including DFEs that arise in mutator strains with differing mutation spectra (Sane et al., 2023; Zeyl and DeVisser, 2001). Empirically-derived fitness landscapes also offer measures of the underlying landscape from which multiple DFEs can be computed (Bank, 2022; Bank et al., 2016; de Visser and Krug, 2014; Szendro et al., 2013).

This availability of data has renewed interest in predicting the shape of the DFE using well-known models such as Fisher’s Geometric model (Martin and Lenormand, 2006, 2015), using fitness landscape theory (Cotto and Day, 2023; Draghi and Plotkin, 2013), or deriving features of the DFE from extreme value theory (Orr, 2003) or at evolutionary equilibrium (Rice et al., 2015). In several of these approaches, the DFE may be predicted for organisms that are either better or more poorly adapted to their environment. While key features of the beneficial tail of the DFE may be independent of the wild type’s state in the adaptive process (Orr, 2003), in general the DFE depends on the fitness of the organism and its adaptive history (Draghi and Plotkin, 2013; Rice et al., 2015).

Since the DFE is predicted to differ for better- or worse-adapted organisms, clearly the DFE must change dynamically during the process of adaptation, a fact highlighted in recent work (Sane et al., 2023). Specifically, if we consider a population adapting to selective pressure, the process of adaptation itself depletes available beneficial mutations, while simultaneously increasing the number of available deleterious mutations (Orr, 2002; Rice et al., 2015; Rokyta et al., 2006). This process re-shapes the DFE as the population adapts. Changes to the DFE as adaptation proceeds have been previously explored in computational simulations (Draghi and Plotkin, 2013), but analytical predictions for changes in the DFE during an adaptive walk have not been developed.

Due in part to this lack of an analytical description, the mechanisms by which the DFE changes as adaptation proceeds are not fully understood. For example, one might expect that when a population is close to a fitness peak, the depletion of the beneficial fraction of the DFE occurs because the availability of high fitness alleles, accessible by mutation, has been considerably reduced. A high fitness allele, however, does not produce a large beneficial effect if it replaces an allele that already provides a good fit to the environment; a beneficial fitness effect also requires that the allele to be replaced stands to be improved. Our approach allows us to disentangle these two effects.

In brief, we study changes to the DFE using a well-studied model of adaptation: a simple adaptive walk (Blanquart et al., 2014; Gillespie, 1983; Kimura, 1991; McCandlish and Stoltzfus, 2014; Orr, 2003) across additive and multiplicative fitness landscapes, in the absence of epistasis. First, we derive analytical approximations for the DFE and the underlying distributions for the relevant fitness contributions, and demonstrate how and by which means the DFE changes as adaptation unfolds. These results are then compared to independent simulations of the adaptive process in which we are able to relax several simplifying assumptions used in the analysis. Our approach offers analytical approximations, at any point on the adaptive walk, not only for the DFE, but also all the underlying distributions of fitness contributions, including the fitness contribution of the next mutation that will fix in the population (Orr, 2003) and the expected fitness over time.

## 2 Methods

We consider a population of haploid individuals whose fitness is characterized by a genotype that encodes *N fitness components*, where each fitness component corresponds to an idealized trait that interacts non-epistatically (but see Discussion) with other traits to determine organismal fitness. Each fitness component can take one of *B* possible alleles (for convenience, we will use *A* to denote the number of alleles available through mutation for each component, such that *A* = *B* − 1). We assume that there exists an underlying fitness landscape, such that each fitness component contributes independently to the overall fitness of the individual by an amount drawn from a distribution of *fitness contributions*, described in the following section. We will consider two models: an additive fitness landscape that is equivalent to both the well-studied NK fitness landscape (Kauffman and Weinberger, 1989) and Orr’s block model (Orr, 2006) (for their non-epistatic cases), and a multiplicative fitness landscape similar to the one described by Nagel et al. (2012).

We model the process of adaptation as an adaptive walk (Gillespie, 1983; Kauffman and Weinberger, 1989; McCandlish and Stoltzfus, 2014). We therefore consider populations in the strong selection-weak mutation regime (Gillespie, 1983): initially all individuals in the population are genetically identical, and when a *de novo* mutation occurs (changing the allele of one fitness component), either all the individuals in the resulting lineage die out, or this lineage spreads to the entire population (it fixes), before another new mutation arises.

### 2.1 Analysis

In this section, we will analyze the sequence of adaptive steps across a fitness landscape. We assume a genotypic, non-epistatic fitness landscape, such that each one of *N* fitness components of the genotype makes a fitness contribution (fc) to the total fitness of the organism. Since each genotype is characterized by the sequence of *N* alleles it encodes for the *N* fitness components, for convenience we will also use “sites” to refer to the loci encoding these fitness components. However we emphasize that these components could be very large-scale traits contributing to fitness (survival, fecundity, growth rate), rather than specific nucleotide sequences.

We will consider both an additive and a multiplicative fitness landscape. For the additive landscape, the total fitness is defined as the average fc across all sites,

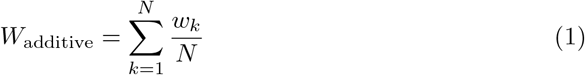

where *w*_*k*_ is the fc of the allele present at the *k*th site of the genotype. On the multiplicative landscape, the total fitness is defined in a similar fashion:

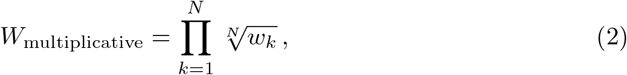

noting that the logarithm of the total fitness in the multiplicative landscape is the average of the log(fc) across all sites.

To develop a stochastic model for these fitness landscapes, we interpret *w*_*k*_ as the random variable for the fc of the allele present at the *k*th site of the initial (or focal) genotype. All sites share an initial probability density function (pdf), denoted as *L*_0_(*w*), where the subscript indicates that zero steps have been taken in the adaptive walk. In this context, Equation 1 defines fitness as the random variable for the mean of *N* independent and identical distributed random variables, each distributed as *L*_0_(*w*) (while Equation 2 defines fitness identically, in log space).

Likewise, let *w*^′^ denote the random variable for the fcs of all the alleles available through mutation (i.e. not present in the focal genotype); these share an initial pdf denoted as *M*_0_(*w*^′^).

An example of the adaptive process is shown in Figure 1, where for illustration we assume each site corresponds to a base pair in the genome. The fcs of alleles in the current genotype, after the *i*th step in the walk, have probability density *L*_*i*_(*w*). If a mutation occurs (where the fc of new mutations has density *M*_*i*_(*w*^′^)), its lineage can either go extinct or fix in the population. If a mutation fixes, the fc of the original and newly fixed alleles on the mutated site may be distributed differently than those on other sites at which a substitution did not occur. We therefore denote the density functions for the original and newly fixed alleles as *L*_*i*|*F*_ (*w*) and *M*_*i*|*F*_ (*w*^′^) respectively. For example, during the first step in the walk, the fc of alleles that might be replaced are drawn from the distribution *L*_0_. Over many replicate walks, the fcs of those alleles that *are* replaced during the first step in the walk are distributed as *L*_0|*F*_.

**Figure 1:**
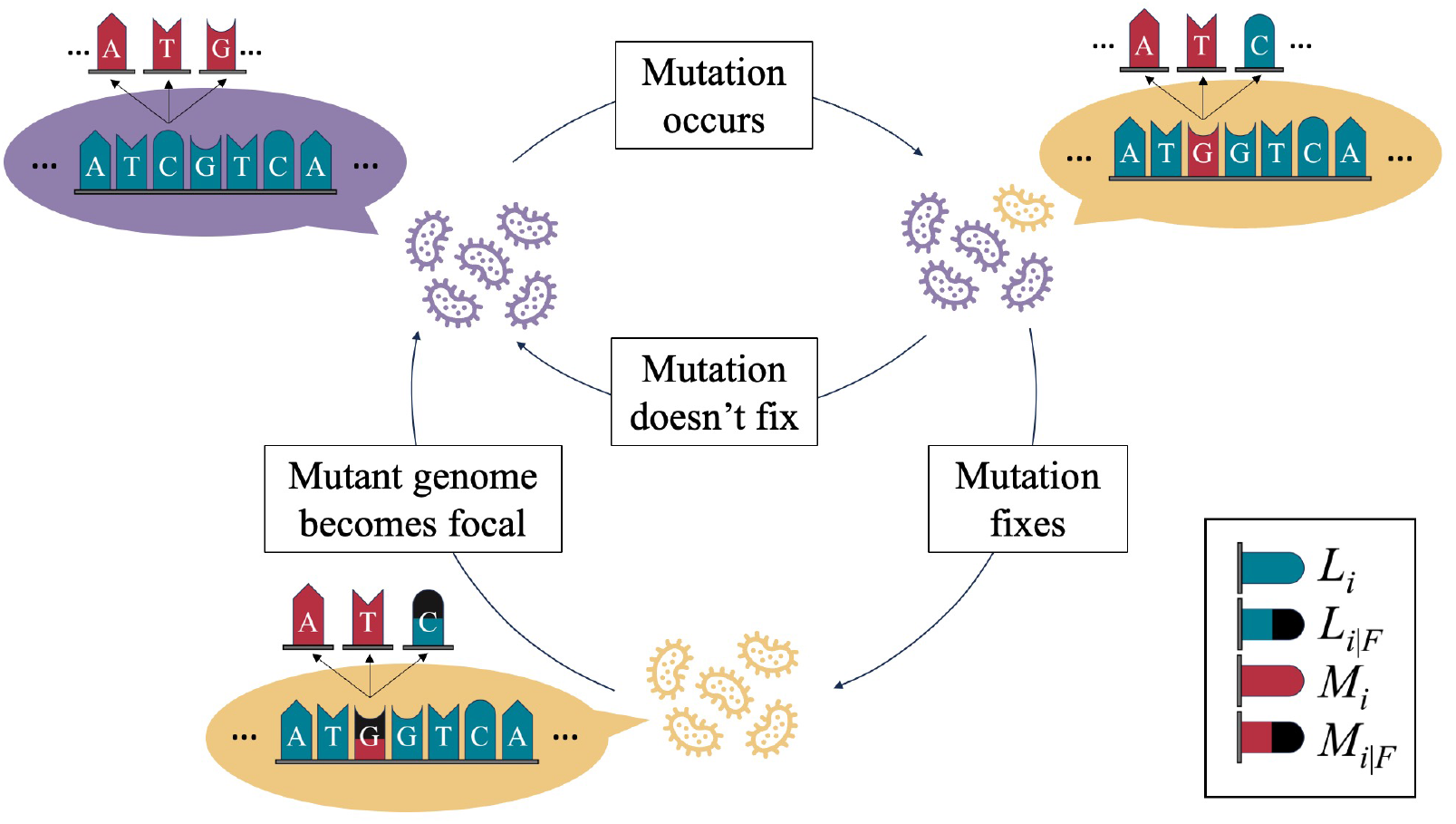
Diagram of the mathematical model after *i* mutations have fixed. Here, we consider a population of clonal haploid organisms (purple) where each fitness component in the genotype correspond to a base pair in the organismal genome. Each fc of each site is drawn from distribution *L*_*i*_ (blue). When a mutant individual (yellow) is produced, the fc of the single mutated site is drawn from *M*_*i*_ (red). The distribution *L*_*i*|*F*_ (blue and black) describes the fc of alleles in the original genotype that were replaced in a fixation event, while *M*_*i*|*F*_ (red and black) describes the fc of alleles that did this replacing (fixed). Once a mutation is fixed, the mutant genotype is considered the new focal genotype, and the process repeats.

It’s also relevant to define the fitness effect, *s*, of a randomly chosen available mutation: if a mutation on the *j*th site would change its fc from *w*_*j*_ = *w* to *w*_*j*_ = *w*^′^, changing the overall fitness of the individual from *W* to *W* ^′^, the fitness effect *s* of that available mutation is defined as

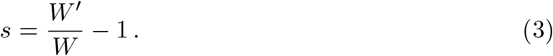

The fitness effect can also be written in terms of *w* and *w*^′^, but its representation will differ for the additive and multiplicative landscapes, respectively:

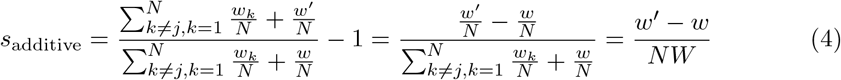

and

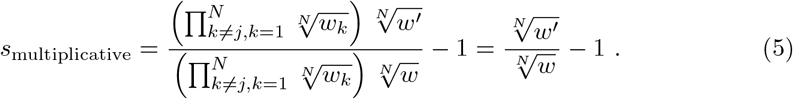

Once a new mutation occurs, that new lineage fixes in the population with a probability of fixation that depends on the fitness effect of the mutation. Letting *F* denote the event that a mutant lineage fixes, we can compute the probability of fixation, Π_0_, of a random mutation using the rule of total probability and conditional probability properties:

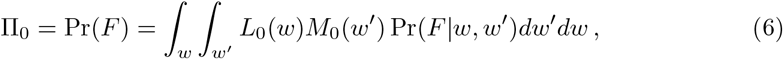

where Pr(*F* |*w, w*^′^) is the probability of fixation of a mutation that changes the fc of the site from *w* to *w*^′^. Here, we use the branching process approximation in Haldane (1927) for small effect beneficial mutations in a large population:

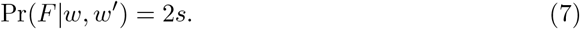

Consider the first step in an adaptive walk, over many replicate trials. Because fixation depends on the fc of the mutation that fixes (*w*^′^), it is clear that the pdf describing mutations that fix in the first step will differ from *M*_0_(*w*^′^), the pdf of all available mutations before the first step. Similarly, the pdf describing the initial fc of all sites, *L*_0_(*w*), differs from the pdf of sites at which a fixation occurs in the first step of the walk, because alleles with lower fc are more likely to be replaced. As described above (see also Figure 1), *L*_0|*F*_ (*w*) and *M*_0|*F*_ (*w*^′^) denote the pdfs of the fc of the site at which the first fixation occurs, for the original and newly fixed alleles, respectively.

We now demonstrate how deriving the two distributions *L*_0|*F*_ and *M*_0|*F*_ will allow us to compute *L*_1_ and *M*_1_. The two distributions *L*_0|*F*_ and *M*_0|*F*_ can be computed using Bayes’ rule for density functions:

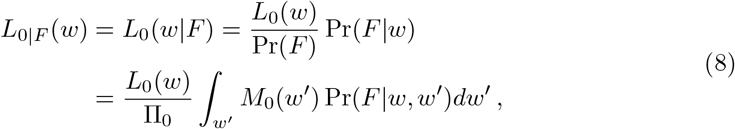

and similarly

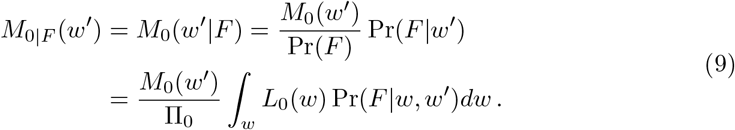

Given these expressions, we can compute the pdf for the fc of the alleles in the remaining *N* − 1 unmutated sites, denoted *L*_0|*U*_, as follows: before the fixation, there were *N* sites each with pdf *L*_0_(*w*), however after the fixation, there is one mutated site with pdf *L*_0|*F*_ (*w*), and *N* − 1 unmutated sites with an unknown pdf (namely *L*_0|*U*_ (*w*)). Since these two sets of alleles are identical, we have

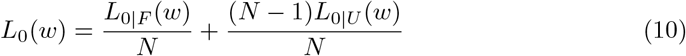

After the fixation event, the mutated genotype is considered the new focal genotype. Therefore, the pdf for the fc of current alleles in the genotype after one step of the walk (denoted *L*_1_(*w*)), takes into account that the fc of a single site has changed its pdf from *L*_0|*F*_ (*w*) to *M*_0|*F*_ (*w*):

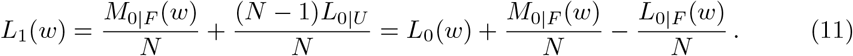

A similar argument can be made to compute the pdf of the available mutations after one step has been taken in the walk, denoted *M*_1_(*w*^′^), with the only difference that now there are *AN* possible alleles to choose as a mutation. The distribution is then given by

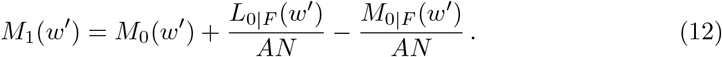

Conveniently, the previous set of equations can be iterated to obtain the corresponding distributions after two mutations have fixed, if we use *L*_1_ and *M*_1_ as initial distributions. In fact, we can iterate this process to obtain the distributions *L*_*i*_ and *M*_*i*_ after *i* mutations have fixed.

### 2.2 Distribution of fitness effects

The distribution of fitness effects (DFE) is defined as the pdf for the fitness effects of all possible available mutations. In this section we will compute the DFE at all steps in the walk, for both the additive and multiplicative fitness landscapes.

On the additive fitness landscape as described in Section 2.1, we can compute the initial DFE, denoted *ρ*_0_(*s*), knowing that the pdf of the sum of two independent random variables is the convolution of their individual distributions. In particular, from Equation 4 we see that a mutation will have fitness effect *s* if the fc of the mutating site changes from *w* before the mutation to *w*^′^ = *w* + *NWs*. This approach is complicated, however, by the fact that *w*, the fitness contribution of a particular site in the genome before mutation, and *W*, the overall fitness of the organism, are not independent. To gain tractability, we make the simplifying assumption that *W* does not depend on *w*, and instead at any step in the walk, the fitness *W* is taken to be a constant equal to its expected value (referred to in the following sections as Assumption 1). The effects of this assumption are discussed in detail in Section S.3, but we note that the assumption should have a negligible effect as long as the number of fitness components, *N*, is large.

Given Assumption 1, and denoting the cumulative density function of *w*^′^ as *C*(*w*^′^), we find that the DFE is given by

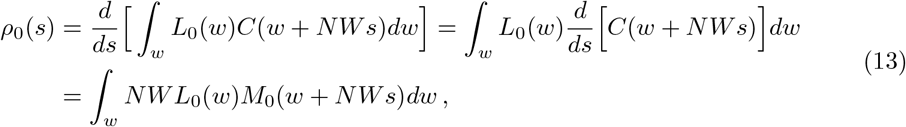

where the limits of integration depend on the support of the fc *w*. In particular, if the fcs have a finite domain, such that *w, w*^′^ ∈ (*α*, Ω), then the the lower limit of integration is given by max(*α, α* − *NWs*) while the upper limit is given by min(Ω, Ω− *NWs*), such that the limits differ for positive and negative *s* values. Although straightforward to calculate, these limits will be suppressed throughout for readability.

Given distributions *L*_*i*_ and *M*_*i*_, we can then follow the approach outlined in Equation 13 to compute the DFE after *i* mutations have fixed. This approach, however, assumes that the fc at any site (distributed as *L*_*i*_) is independent of the fcs of the available mutations at that site (distributed as *M*_*i*_). This assumption does not hold for sites at which a mutation has occurred, since the previously replaced allele is now part of the set of available mutations. To proceed, we make a second simplifying assumption: that the fc at any site is independent of the fc of available mutations at that site (referred to as Assumption 2). The implications of this assumption are examined further in Section S.3.

Under Assumption 2, the DFE after *i* mutations have fixed is given by

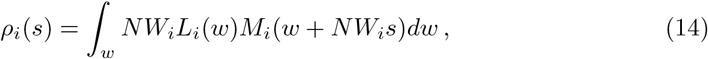

where again *W*_*i*_ is the expected fitness after *i* mutations have fixed (Assumption 1), and the limits of integration are defined as before.

On the multiplicative fitness landscape, we can follow a similar process to compute the initial DFE, denoted *σ*_0_(*s*). In this case, from Equation 2 we can see that a mutation will have fitness effect *s* if the fc of the mutating site changes from *w* before the mutation to *w*^′^ = *w*(1 + *s*)^*N*^. Given that the random variables *w* and *w*^′^ are initially independent, the DFE is given by the convolution of the pdf of these variables:

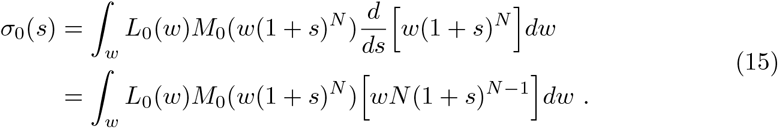

Similarly, given the corresponding distributions *L*_*i*_ and *M*_*i*_, we can follow the previous approach to compute the DFE after *i* mutations have fixed. As before, *w* and *w*^′^ are not independent in all sites after *i* mutations have occurred, therefore Assumption 2 is imposed in the multiplicative case as well. Then, the DFE after *i* mutations have fixed is

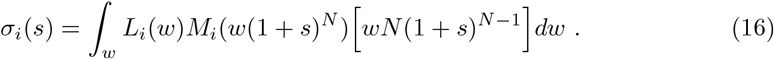

### 2.3 Implementation of the equations

Given an initial distribution for the fitness contributions (we illustrate several in the results to follow) we used Maple (Maplesoft, 2022) to exactly compute *L*_*i*_, *L*_*i*|*F*_, *M*_*i*_ and *M*_*i*|*F*_, iterating until we reached a given number of mutations. Unfortunately, computational times increase exponentially with the number of mutations. For example, in the simplest case considered (on an additive fitness landscape when *L*_0_ = *M*_0_ = *U* (0, 1), the uniform distribution on (0,1)), the distributions *L*_*i*_ and *M*_*i*_ are polynomials of degree 2^*i*+1^ − 2, which translates to computational times that grow in a similar fashion.

We therefore also compute the equations numerically using Python, which reduces the computational times significantly (and eliminates the exponential dependence on the number of steps taken). In this case, the distributions *L*_*i*_, *L*_*i*|*F*_, *M*_*i*_ and *M*_*i*|*F*_ as well as the DFEs were computed numerically as vectors of length 10^4^. As a test case, we computed the error of the numerical calculations with respect to the exact calculations for *N* = 20, *m*_max_ = 5 and *L*_0_ = *M*_0_ = *U* (0, 1); the largest absolute error in the DFE in this case was ≤ 10^−4^. Finally, for all the results shown here, numerical integrations were computed with absolute and relative errors bellow 10^−4^ for the fc distributions and 10^−3^ for the DFEs.

### 2.4 Simulations

To test the accuracy of our analytical approximations, we performed computer simulations of adaptive walks across non-epistatic additive and multiplicative fitness landscapes. Here, we considered the number of alleles per site to be *B* = 2, 4, 20 as well as the limiting case of an infinite number of alleles per site.

For each adaptive walk, we draw the fitness contributions of each allele on each of the *N* sites of the genotype independently from a specified distribution. We show the results for the case in which the fcs are drawn from uniform distributions: for the additive fitness landscape, we considered a uniform distribution with range (0, 1), while for the multiplicative fitness landscape we considered a range (0.5, 1.5), such that fcs are centered around unity.

Throughout the results shown, the allele at each site is chosen at random to form the initial sequence. While this initial condition is clearly at odds with reality for an adapting population, in the SI (Section S.2) we explore the case in which alleles are chosen such that the current genome is relatively “fit” initially. We also point out that our approach holds if we take *N* to be some number of fitness components that are not optimized (after an environmental shift, for example), whereas other components of the genome will remain unchanged in the adaptive process under study.

Starting from an initial sequence where the allele on each site is chosen at random, a mutant sequence is created by choosing a site uniformly at random, and changing the allele at that site uniformly at random to one of the *B* − 1 other alleles. The probability of fixation of the mutation is then computed according to Equation 7, and the fate of the mutation is randomly chosen accordingly. If the mutation fixes, the mutant sequence becomes the new focal sequence, and a step in the walk is taken. Otherwise, the focal sequence remains the same, a new mutation is proposed, and the process repeats.

The adaptive walk ends when the number of fixed mutations reaches a desired value. At any point in the walk, we can examine the fcs of the alleles in the current focal sequence, as well as the fcs of all available mutations, which we use to generate DFEs. Simulation results were obtained by simulating 10^5^ adaptive walks, each over a different landscape.

We note that these simulations relax both Assumptions 1 and 2 from the analytical work. First, in the simulations the fitness of the current sequence, *W*, is not independent of the fc of any particular site, *w*_*i*_, and rather than taking a mean value, it differs for each adaptive walk. Second, the DFEs computed in the simulations compute each *s* value by comparing the fc of each site with the *B* − 1 available mutations at that site, therefore the dependency at substituted sites is preserved.

## 3 Results

First, we compare the analytical approximations and simulation results for the four key distributions in our analysis: *L*_*i*_, *L*_*i*|*F*_, *M*_*i*_ and *M*_*i*|*F*_, as the adaptive walk progresses. Later, we study the dynamics of the DFE as the adaptive walk progresses. We considered additive and multiplicative fitness landscapes, although most results shown in this section will be from additive fitness landscapes, as results are very similar for both models (see Section 4). Further results can be found in Section S.1 as indicated below.

Unless stated otherwise, results are shown for phenotypes of length *N* = 100, and considering four parameter values for the number of possible alleles for each fitness component: *B* = 2, 4, 20 and the limiting case in which there are infinite possible alleles. As mentioned before, initial uniform distributions were assumed for the fitness contributions in the both landscapes. For comparison across cases, adaptive walks were performed until the beneficial fraction of available mutations in the simulations fell bellow 15%.

### 3.1 Fitness contributions

In Figure 2, we show the the initial distributions for *L*_*i*_, *L*_*i*|*F*_, *M*_*i*_ and *M*_*i*|*F*_, comparing each with the same distribution at the end of the adaptive walk, for the additive and multiplicative landscapes, respectively, at *B* = 4. The corresponding figures for the multiplicative fitness landscapes, and *B* = 2, 20 and the infinite alleles model can be seen in Section S.1. Overall we find excellent agreement between the analytical approximation (solid lines) and simulation results (histograms). As adaptation proceeds, the pdf of fcs in the current genotype, *L*_*i*_, is reshaped into a distribution with a higher expected value (shifts right, top left panel); conversely, the pdf of available mutations, *M*_*i*_, is reshaped into a distribution with a lower expected value (shifts left, top right panel). This is expected for an adaptive walk, in which the mutations that fix are more likely to occur at sites where an allele with low fc (bottom left panel) is replaced by an allele with a high fc (bottom right panel).

**Figure 2:**
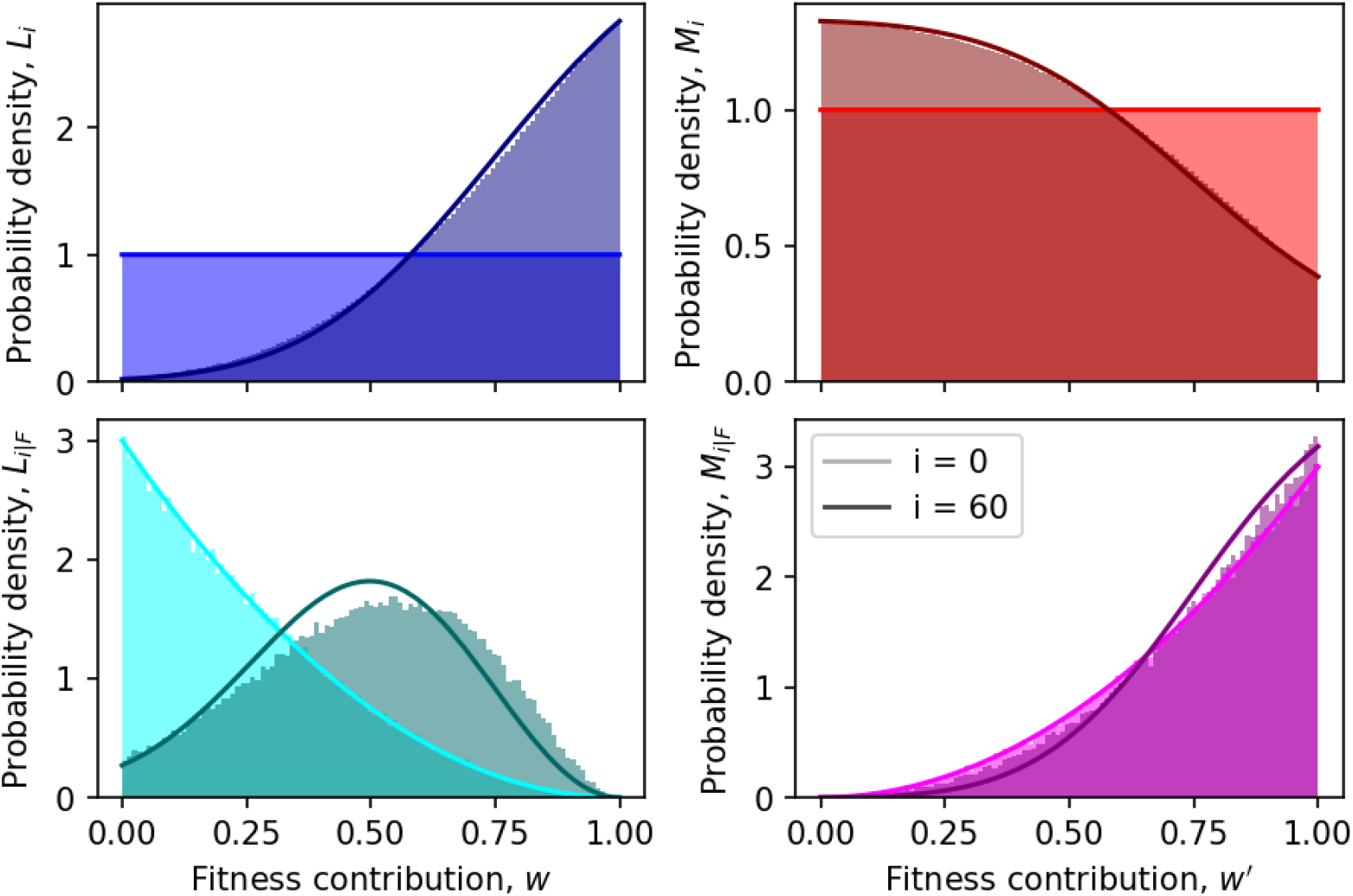
Simulation results (colored histograms) and analytical approximations (solid lines) for the density functions *L*_*i*_ (blue), *L*_*i*|*F*_ (cyan), *M*_*i*_ (red) and *M*_*i*|*F*_ (magenta) in an additive fitness landscape. The initial distributions (at *i* = 0) are in lighter colors, while the distributions for *i* = 60 are in darker colors. Results for *B* = 4, *N* = 100, *L*_0_ = *M*_0_ = *U* (0, 1) are shown.

Interestingly, we find that the distributions of fc in the focal genotype (*L*_*i*_ and *L*_*i*|*F*_) change more dramatically than the distributions of available mutations (*M*_*i*_ and *M*_*i*|*F*_). This presumably arises because for these examples there are *A* = 3 available mutations per site, and therefore each fixation event has a more modest effect on the overall distribution of available mutations compared to its effect on the distribution of focal fcs.

In Figure 3 and we show how the mean value of each of these four distributions changes as the walk progresses, again comparing analytical and simulation results, for the additive and multiplicative landscapes, respectively. Here, the distributions *L*_*i*_ and *M*_*i*_ initially have the same mean, but as mutations accumulate the mean fc of alleles that comprise the focal genotype (blue) increases, while the mean fc of available mutations (red) decreases. For the cases where *B >* 2 there are fewer alleles in the genotype than available through mutation, and thus *L*_*i*_ increases faster than *M*_*i*_ declines. The increase in *L*_*i*_ corresponds to an increase in genotype fitness over time, as expected in adaptive walks (illustrated in Figure S.18). We also see that the mean for *L*_*i*|*F*_ (alleles that are replaced in the *i*th step of the walk) increases. This is reasonable since low fitness alleles are more likely to be replaced early in the walk, and fewer remain as the walk progresses.

**Figure 3:**
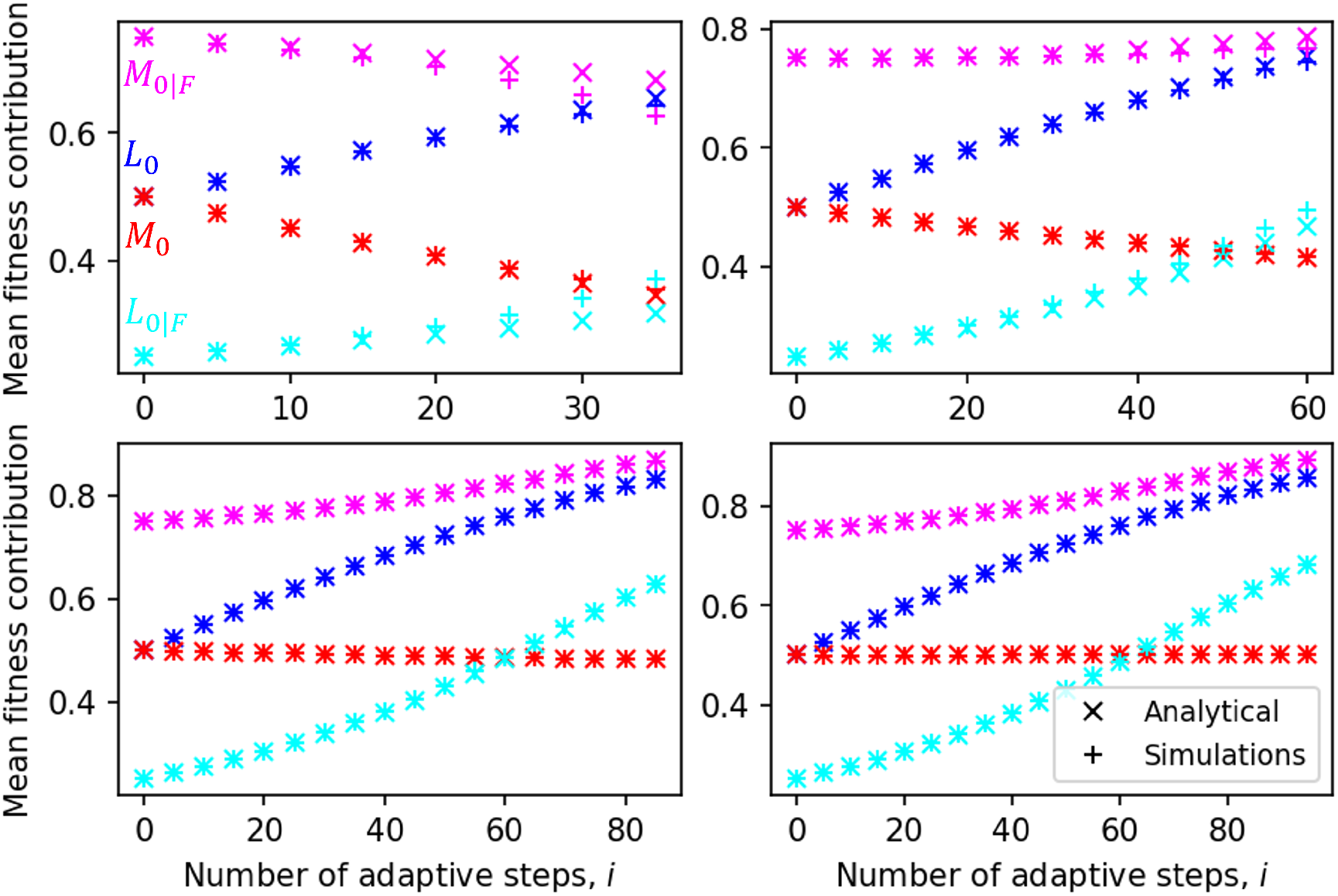
Analytical approximations (crosses) and simulation results (plus signs) for an additive landscape for the four distributions for the fitness contributions: *L*_*i*_ (blue), *L*_*i*|*F*_ (cyan), *M*_*i*_ (red) and *M*_*i*|*F*_ (magenta), for different number of available alleles at each site, *B*. In each case, *i* steps in the adaptive walk are taken such that the beneficial fraction of the DFE falls below 15%. For *B* = 2 (top left), 4 (top right), 20 (bottom left), and the infinite alleles model (bottom right), the numbers of steps taken are *i* = 35, 60, 85 and 95, respectively. Results are shown for *N* = 100 and *L*_0_ = *M*_0_ = *U* (0, 1).

Finally, Figure 3 demonstrates that as adaptation proceeds the mean fc of the mutations that fix, *M*_*i*|*F*_ (purple symbols), is dependent on the number of alleles per site. For sufficiently high *B*, the mean of this distribution increases over time, which implies that adaptation is not limited, in these simulations, by the depletion of available mutations with high fc. Taken together with the increase in the fc of the alleles that are replaced, *L*_*i*|*F*_ (green symbols), this suggest that the availability of low-fitness alleles to replace is a stronger limiting factor for adaptation. We will return to this idea in the following section.

### 3.2 Distribution of fitness effects

Figure 4 shows the analytical predictions for the DFE as adaptation proceeds, for uniform initial distributions of fcs. As mutations accumulate, we see in both landscapes a shift in the probability density towards more negative fitness effects, as a result of the fixation of beneficial mutations. Not only are beneficial mutations depleted by fixation, but each beneficial mutation, once fixed, contributes a negative fitness effect to the DFE. In the additive fitness landscape we see that the DFE gradually narrows over time, such that available fitness effects, on average, have smaller magnitudes. Given the definition of the selective effect (Equation 4), we see that this occurs because the fitness of the focal genotype increases with each fixed mutation.

**Figure 4:**
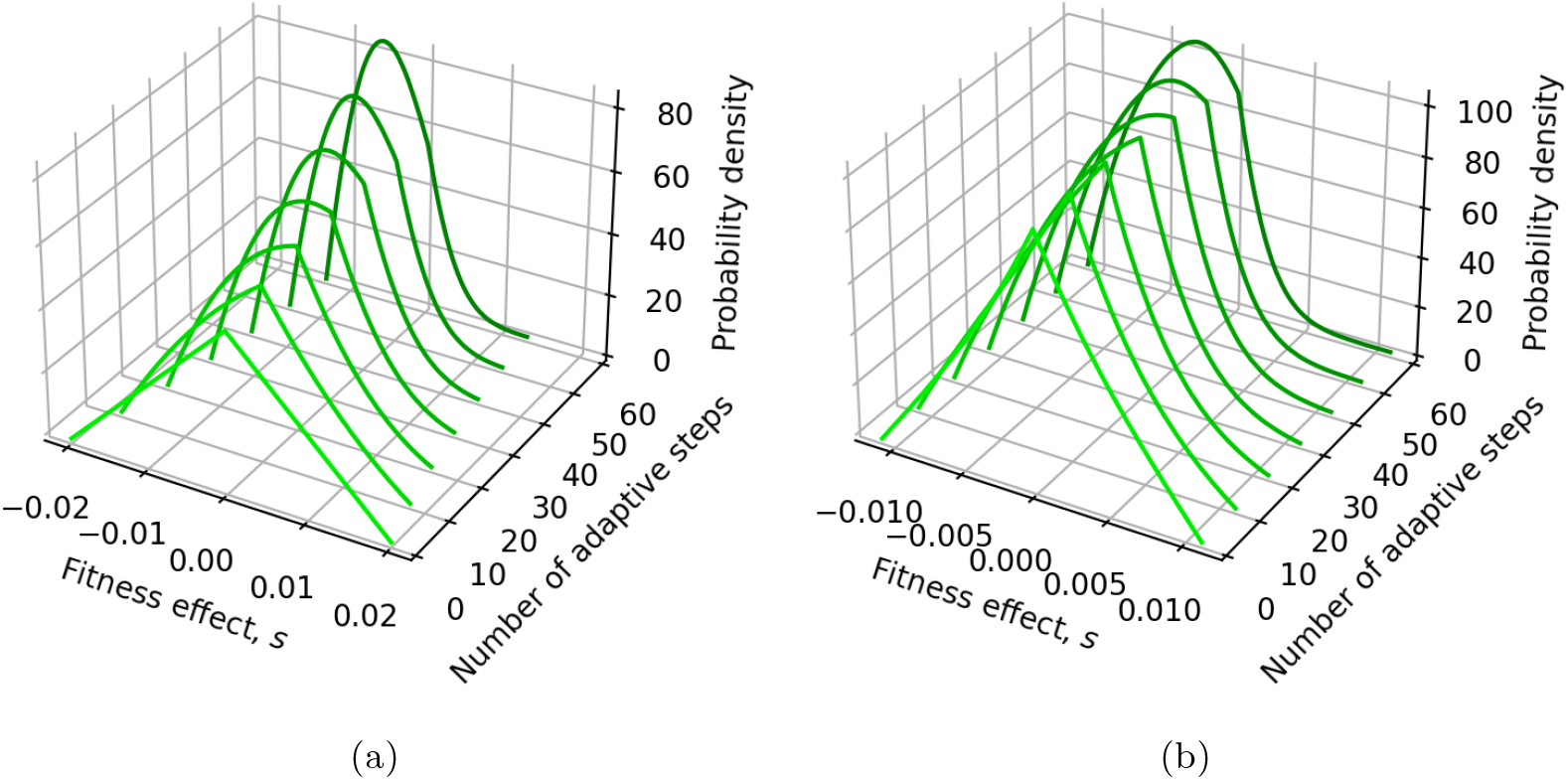
The DFE changes shape as the number of steps in the adaptive walk increases. (a) Analytical results for an additive landscape and initial fc distributions *L*_0_ = *M*_0_ = *U* (0, 1). (b) Analytical results for a multiplicative landscape and initial fc distributions *L*_0_ = *M*_0_ = *U* (0.5, 1.5). Results are shown for *N* = 100 and *B* = 4.

In Figures 5 and 6 we show a comparison between analytical approximations and simulation results for the DFEs and distributions *L*_*i*_ and *M*_*i*_, at the end of the adaptive walk. Overall, the analytical predictions for the DFEs capture the qualitative behaviour well, and become increasingly accurate as *B* increases. As discussed further in the SI (Section S.3), discrepancies at *B* = 2 and 4 arise from Assumption 2, that *w* and *w*^′^ are independent random variables. In reality, if a mutation has previously fixed at a specific site, that site tends to have a higher fc, while one of the *B* − 1 available mutations at that site corresponds to the allele that was replaced, which tends to have a lower fc; this creates a dependency between the random variables *w* and *w*^′^ at those sites.

**Figure 5:**
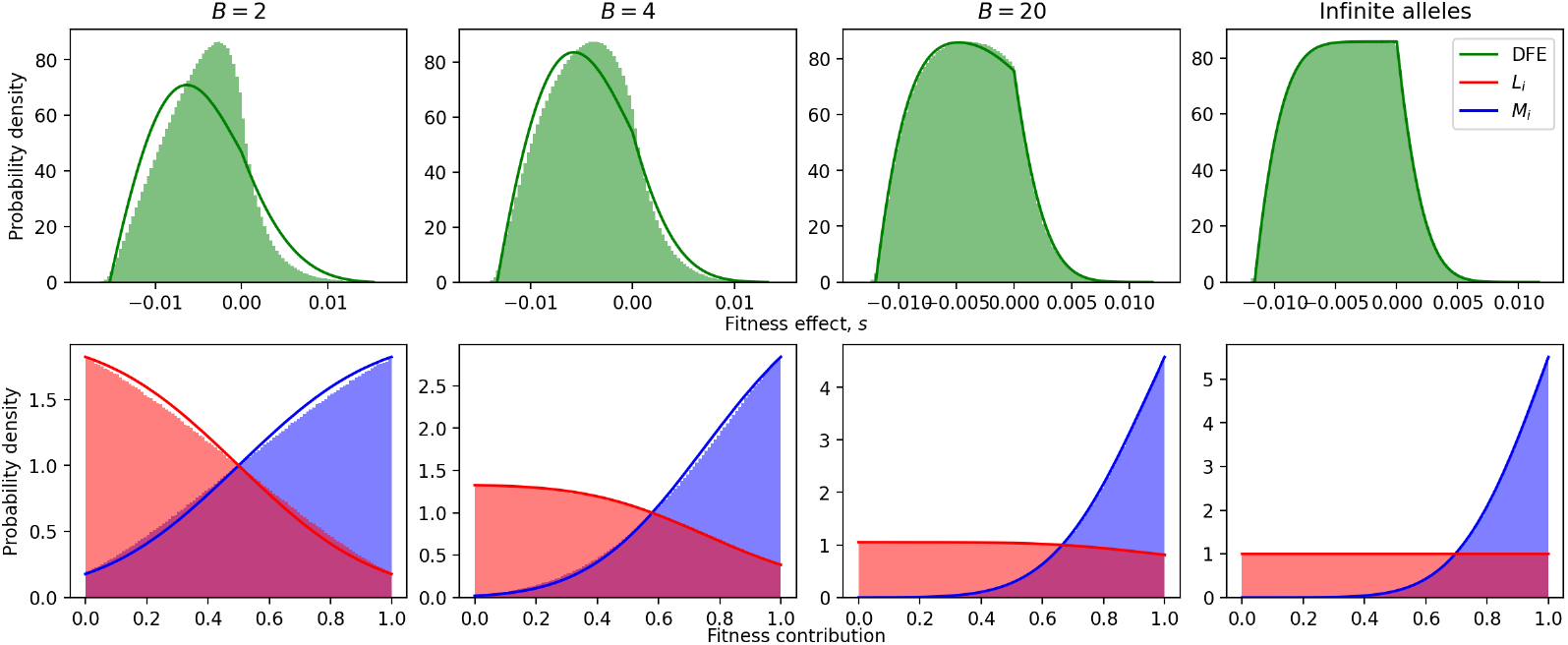
Analytical approximations (solid lines) and simulation results (histograms) for the DFE (top row) and distributions *L*_*i*_ and *M*_*i*_ (bottom row), for different number of available alleles at each site, *B*, for an additive fitness landscape. In each case, the number of steps in the adaptive walk are such that the beneficial fraction of the DFE falls just below 15%. For *B*=2, 4, 20 and the infinite alleles model, the numbers of steps taken are 35, 60, 85 and 95, respectively. Despite the near-constant beneficial fraction of the DFE, the distributions of available fitness contributions differ significantly across cases. Results are shown for *N* = 100 and *L*_0_ = *M*_0_ = *U* (0, 1).

**Figure 6:**
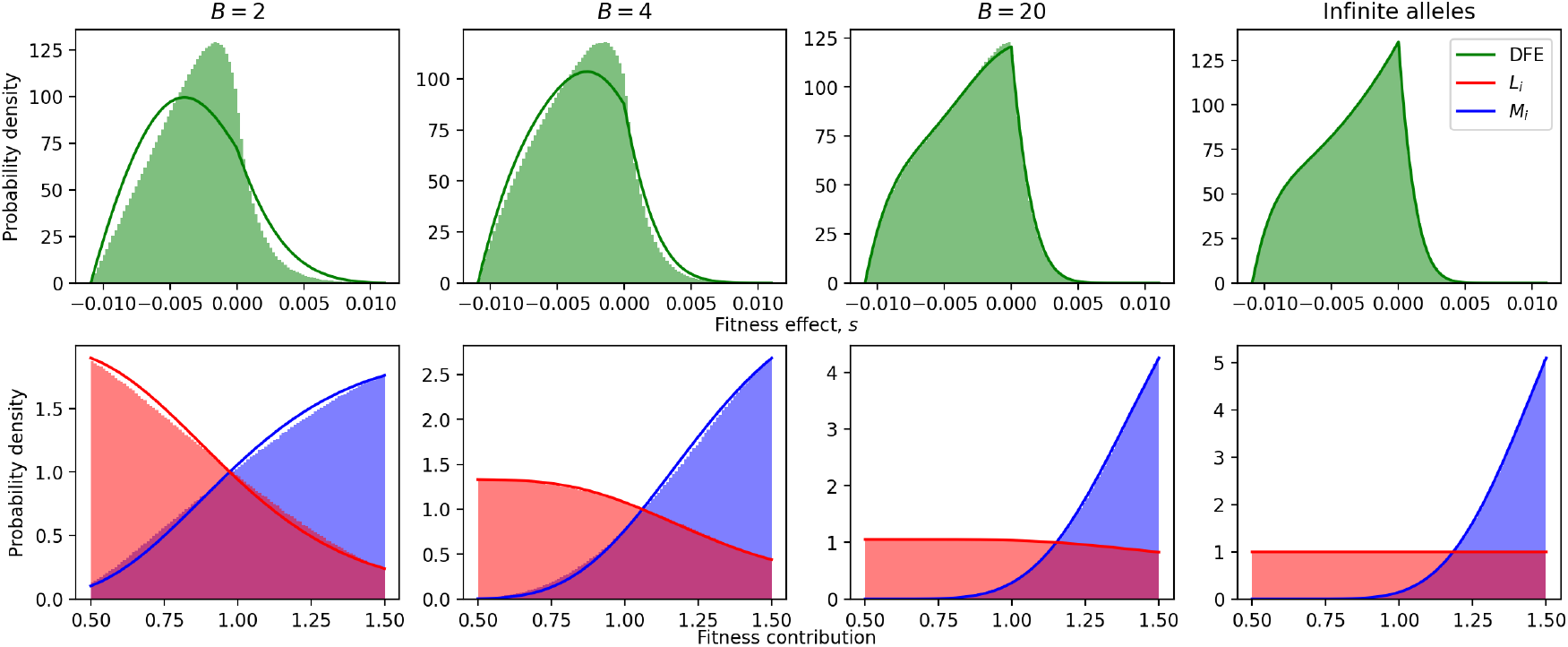
Analytical approximations (solid lines) and simulation results (histograms) for the DFE (top row) and distributions *L*_*i*_ and *M*_*i*_ (bottom row), for different number of available alleles at each site, *B*, for a multiplicative fitness landscape. In each case, the number of steps in the adaptive walk are such that the beneficial fraction of the DFE falls just below 15%. For *B*=2, 4, 20 and the infinite alleles model, the numbers of steps taken are 35, 60, 85 and 95, respectively. Despite the near-constant beneficial fraction of the DFE, the distributions of available fitness contributions differ significantly across cases. Results are shown for *N* = 100 and *L*_0_ = *M*_0_ = *U* (0.5, 1.5).

As seen in Figures 5 and 6, despite near-identical beneficial fractions in the DFE, the distributions *L*_*i*_ and *M*_*i*_ are markedly different when there are *B* = 2, 4, 20 or an infinite number of alleles available at each site. In particular, when *B* is large, the distribution of available fitness contributions *M*_*i*_ differs little from the initial uniform distribution, while the distribution *L*_*i*_ changes drastically. We conclude that the 15% beneficial fraction in the DFE is achieved via different mechanisms in each case. In particular, at *B* = 20 it is clear that the beneficial fraction of the DFE is limited by the depletion of sites in the genotype currently carrying low-fitness alleles (left-most part of blue distributions), rather than by the availability of high-fitness mutations (right-most part of red distributions). This effect is also apparent, though more subtle, when *B* = 4, whereas when *B* = 2 the beneficial fraction of the DFE is equally limited by current low-fitness alleles and available high-fitness mutations.

## 4 Discussion

Previous analytical descriptions of the distributions of fitness effects have demonstrated that the DFE differs, as expected, for individuals that are better-or worse-adapted to the environment (Cotto and Day, 2023; Martin and Lenormand, 2006, 2015; Orr, 2003). Extending these ideas, Rice et al. (2015) treated the DFE as a dynamic distribution, and used substitution rates of beneficial and deleterious mutations to predict the shape of the DFE at evolutionary equilibrium. Interestingly, simulated equilibrium DFEs in that approach are very similar to the results we obtain after longer periods of adaptation (see Figure 2E in Rice et al. (2015) for comparison).

Although fixed or equilibrium DFEs have thus been well studied, recent experimental and theoretical work has highlighted the fact that the DFE can change significantly during adaptation (Sane et al., 2023). Our aim was to give a quantitative description of the process of adaptation to describe the underlying forces that change the DFE as a population adapts.

Using a highly simplified and thus tractable model of the adaptive process, we provide analytical expressions for the DFE, at any step in an adaptive walk, given an additive or a multiplicative fitness landscape and an arbitrary distribution of independent fitness contributions. Consistent with intuitive expectations, we demonstrate that as adaptation proceeds, the DFE is reshaped in such way that the beneficial fraction decreases (and therefore the deleterious fraction increases). This increase in the fraction of deleterious mutations occurs because mutations that fix tend to be beneficial, and once fixed each of these mutations contributes a corresponding deleterious (reverse) mutation to the DFE (see Rice et al. (2015)). On the additive fitness landscape, we observe a decrease in the variance of the DFE (see Figure 4), which is a consequence of the classic definition of the fitness effect, *s* (Equation 4): as the mean population fitness increases, mutations with the same absolute (additive) fitness contribution yield smaller values of *s*. Notice that this decrease in variance of the DFE is not observed if a multiplicative fitness landscape is assumed, since the fitness effect in this model is independent of the population fitness (Equation 5).

Although the DFE shows similar dynamics when the number of alleles per fitness component (parameter *B*) is varied, the underlying distributions of fitness contributions differ markedly across the studied cases. We examined this trend in Figures 5 and 6, where we compared adaptive walks that reached the same total depletion of the beneficial DFE, but occurred on landscapes with different numbers of available alleles per site. Here we observed that as the number of alleles per site increases, the mechanism underlying the overall depletion of the DFE shifts. When *B*=2, the beneficial fraction of the DFE is equally affected by two factors: the depletion of low-fitness alleles in the focal genome that can be “improved” by a substitution, and the depletion of high-fitness available mutations. As long as *B* exceeds two, however, these two factors no longer balance, and in fact the depletion of low-fitness alleles in the focal genotype becomes the more important limiting factor. This implies that adaptation, or at least the idealized adaptive process studied here, is less limited by the availability of beneficial mutations, and more limited by the availability of low-fitness alleles to be replaced.

How likely is this prediction to be relevant in nature? The fixation probability of a beneficial mutation depends on the change in fitness achieved by the new mutation. If we assume that fitness is additive or multiplicative at some scale (possibly the additive or multiplicative contributions of proteins, operons or larger networks), then the fixation probability will depend with equal weight on the fitness contribution of the new, mutated operon (say) and on the fitness contribution of the existing operon. Suppose there are *N* operons in the current genome, and *A* ways in which each operon could be mutated. We see that *A* is possibly a very large number, and thus the total number of available mutations, *AN*, far exceeds the number of fitness components, *N*, in the genotype. We can thus imagine a situation in which there are many available beneficial mutations, but they only achieve a non-negligible fixation probability if they improve the fitness of a comparatively low-fitness operon. Overall, we suggest that when an underlying fitness landscape is additive or multiplicative, adaptive processes may well be limited by the availability of low-fitness components that need improvement, rather than the availability of high-fitness mutations to do the improving. The extent to which this prediction holds, more generally, is a clear avenue for future study.

The results shown here consider non-epistatic fitness landscapes, which simplify the analysis and reduce computational times, but are highly idealized. In particular, we notice that the overall size of the fitness effect of new mutations, *s*, is considerably smaller than empirical measurements of this quantity (Bank et al., 2016; Kassen and Bataillon, 2006; Sane et al., 2023). This occurs because a single point mutation can change the fitness of only one of *N* fitness components; in contrast, on an epistatic landscape each mutation can affect several fitness components, allowing for larger magnitude fitness effects. It is also not surprising that our results for the additive and multiplicative landscapes are similar (see for example Figures 5 and 6), due to the fact that the fixation probability in these models converges when mutations carry small fitness effects, as described by Nagel et al. (2012).

During the adaptive walk, we assume a large population size and thus allow only beneficial mutations to reach fixation. A further extension of this work would allow both beneficial and deleterious substitutions, complicating the DFE dynamics but allowing it to reach an equilibrium as adaptation progresses (Rice et al., 2015).

Our analytical approach for computing the DFE makes several further assumptions which substantially simplify the necessary computation. Despite these simplifications, the approximation performs well, describing the dynamics of the DFE and the underlying distributions on which it depends. After a substantial fraction of the genome has been subject to a substitution event, however, discrepancies between the approximation and simulation results become apparent. In the SI (Section S.3) we demonstrate how these errors are related to the simplifying assumptions in our model, and show that these errors vanish if we relax those assumptions in the analysis or impose them in the simulations. Analytical expressions for the DFE that relax these assumptions (for more than the initial DFE, as calculated in Section S.3) should be achievable in future work.

We also note that these discrepancies are reduced in realistic parameter regimes. Since the variance of the pdf for fitness *W* is inversely proportional to the number of fitness components *N*, fitness *W* will be very close to its mean value when a larger number of components are considered. We also point out that discrepancies emerge when a large fraction of the genome, 20-60%, has been subject to an adaptive substitution. Finally, as we increase the number of alleles available at each site, the dependency of the current fitness contribution and the fitness contribution of a proposed mutation is reduced. Although our simulations were limited for computational tractability, each of these factors would increase the accuracy of the analytical predictions derived here.

We have outlined an analytical approach that allows for a directly computable prediction of a dynamic DFE. While we show results for an idealized adaptive process, this approach could clearly be expanded in future to study an epistatic fitness landscape (Bank, 2022), for example by varying the parameter *K* in the NK model (Kauffman and Weinberger, 1989). Another simplifying assumption in our approach is that all mutations are equally likely to occur. In reality, mutation biases are ubiquitous, across the tree of life and at multiple genomic scales (Couce et al., 2013; Katju and Bergthorsson, 2018; Stoltzfus and Yampolsky, 2009). Applying this approach to predict the DFE during biased adaptation is thus another interesting avenue for future work. More generally, it is our hope that future work in this direction will help close the gap between empirical and theoretical descriptions of the distribution of fitness effects (Bataillon and Bailey, 2014; Cotto and Day, 2023; Kassen and Bataillon, 2006).

## Acknowledgments

This work was supported by the Natural Sciences and Engineering Research Council of Canada grant RGPIN-2019-06294.

## S Supplementary Information

### S.1 Results for alternative numbers of alleles

In this section we show results analogous to those shown in Section 3, for an additive and a multiplicative fitness landscape, for alternative numbers of alleles per site, B. Analogous to Figure 2, for an additive fitness landscape, we have Figures S.1, S.2 and S.3 for *B* = 2, 20 and the infinite alleles model, respectively. For a multiplicative landscape, we have analogous results in Figures S.4, S.6 and S.7. Notice that both models show very similar results despite the differences in their formulation, if we shift the range for the fitness contributions. Analogous to Figure 3, we have Figure S.8 for a multiplicative fitness landscape. Again, the results for both models are very similar. Finally, we have analogous results to Figure 4 for different number of alleles per site, B, in Figures S.9, S.10 and S.11 for *B* = 2, 20 and an infinite alleles model, respectively.

**Figure S.1:**
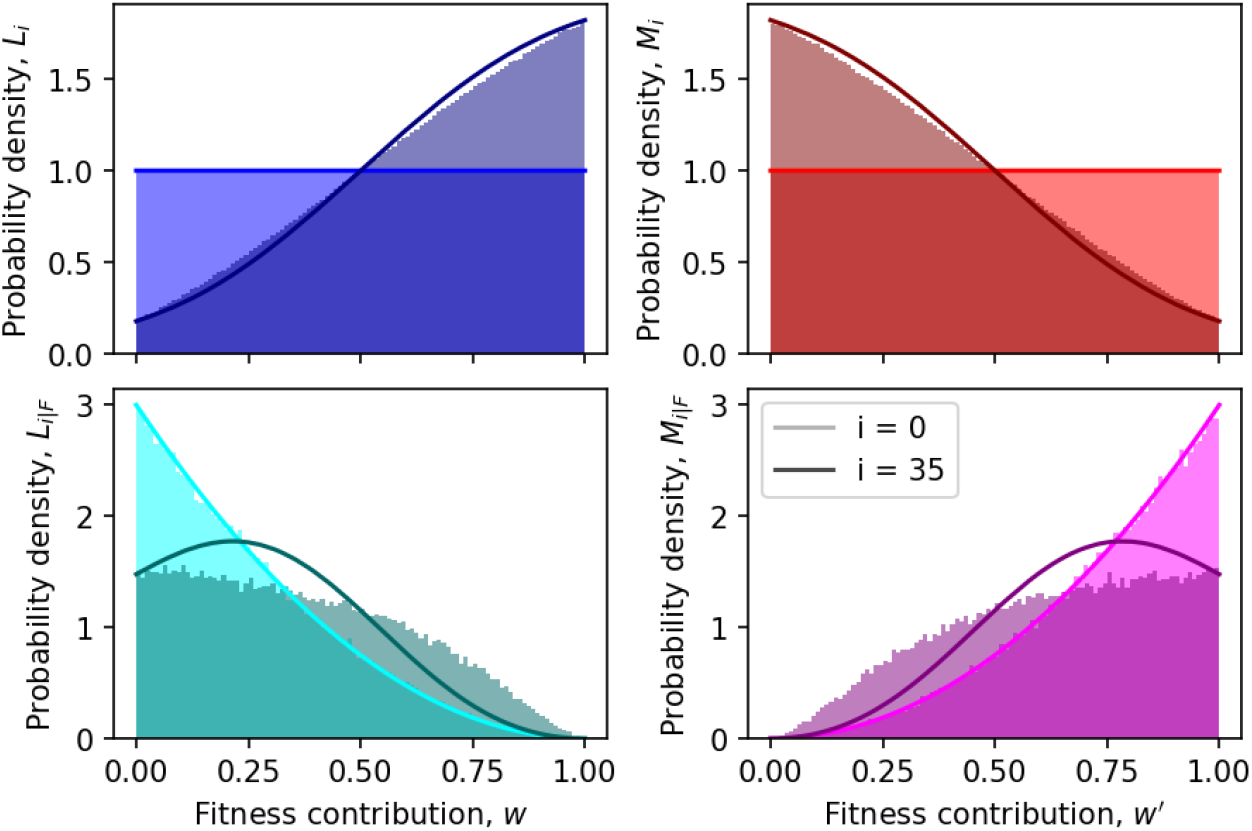
Simulation results (colored histograms) and analytical approximations (solid lines) for the density functions *L*_*i*_ (blue), *L*_*i*|*F*_ (cyan), *M*_*i*_ (red) and *M*_*i*|*F*_ (magenta) in an additive fitness landscape. The initial distributions (at *i* = 0) are in lighter colors, while the distributions for *i* = 35 are in darker colors. Results for *B* = 2, *N* = 100, *L*_0_ = *M*_0_ = *U* (0, 1) are shown.

**Figure S.2:**
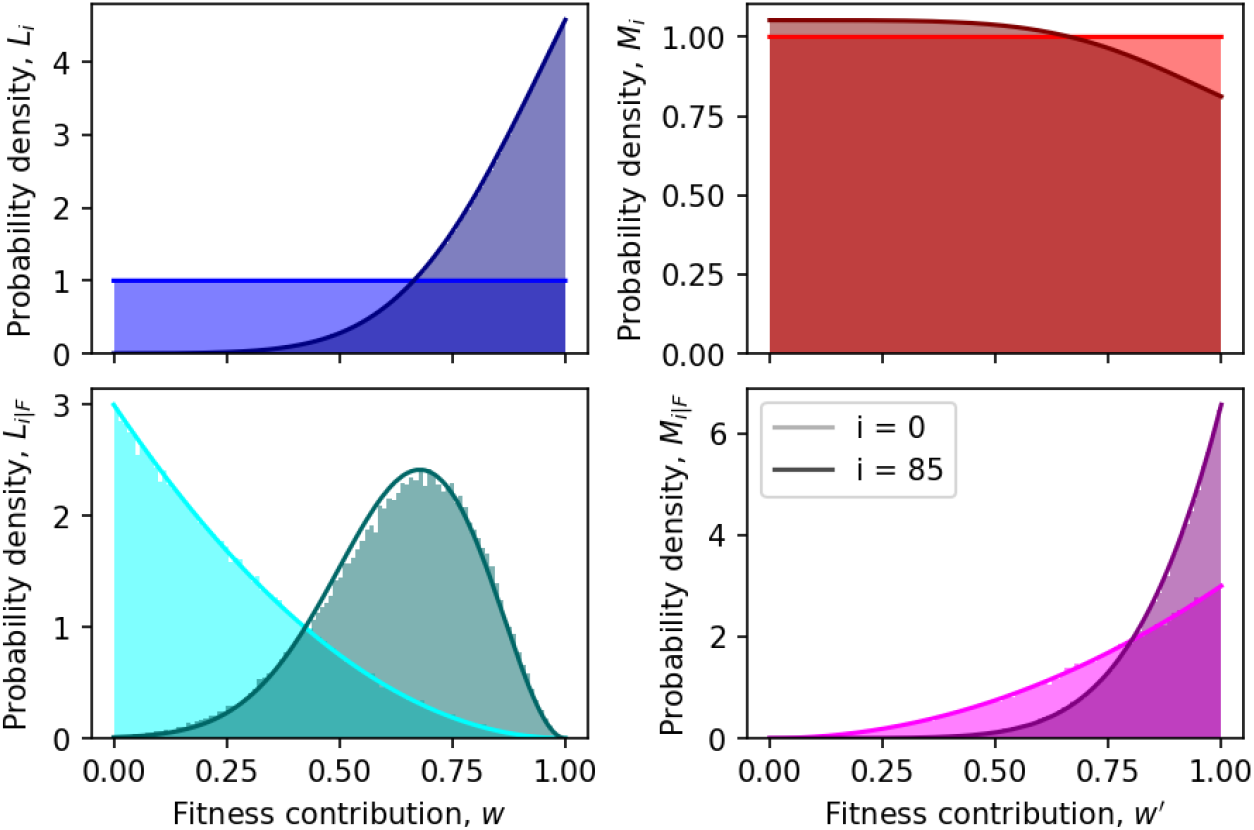
Simulation results (colored histograms) and analytical approximations (solid lines) for the density functions *L*_*i*_ (blue), *L*_*i*|*F*_ (cyan), *M*_*i*_ (red) and *M*_*i*|*F*_ (magenta) in an additive fitness landscape. The initial distributions (at *i* = 0) are in lighter colors, while the distributions for *i* = 85 are in darker colors. Results for *B* = 20, *N* = 100, *L*_0_ = *M*_0_ = *U* (0, 1) are shown.

**Figure S.3:**
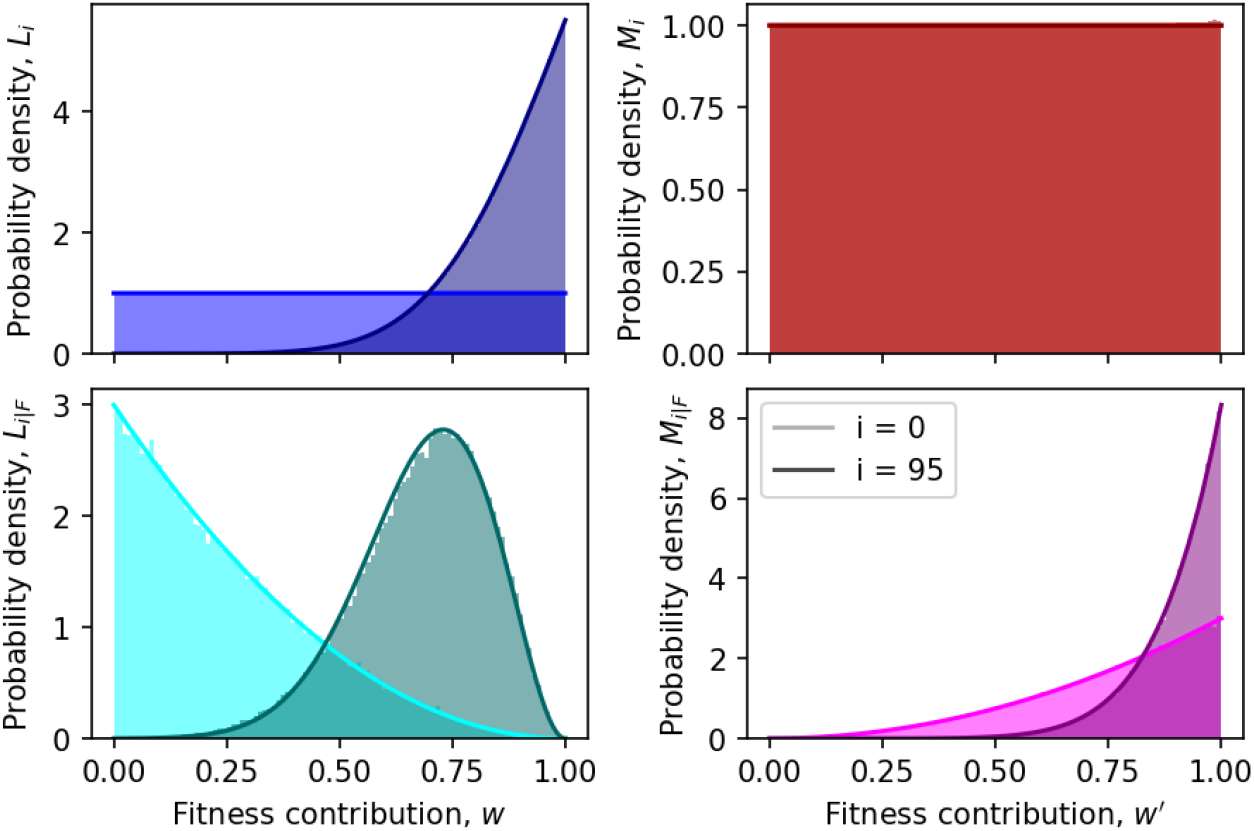
Simulation results (colored histograms) and analytical approximations (solid lines) for the density functions *L*_*i*_ (blue), *L*_*i*|*F*_ (cyan), *M*_*i*_ (red) and *M*_*i*|*F*_ (magenta) in an additive fitness landscape. The initial distributions (at *i* = 0) are in lighter colors, while the distributions for *i* = 95 are in darker colors. Results for an infinite alleles model, *N* = 100, *L*_0_ = *M*_0_ = *U* (0, 1) are shown.

**Figure S.4:**
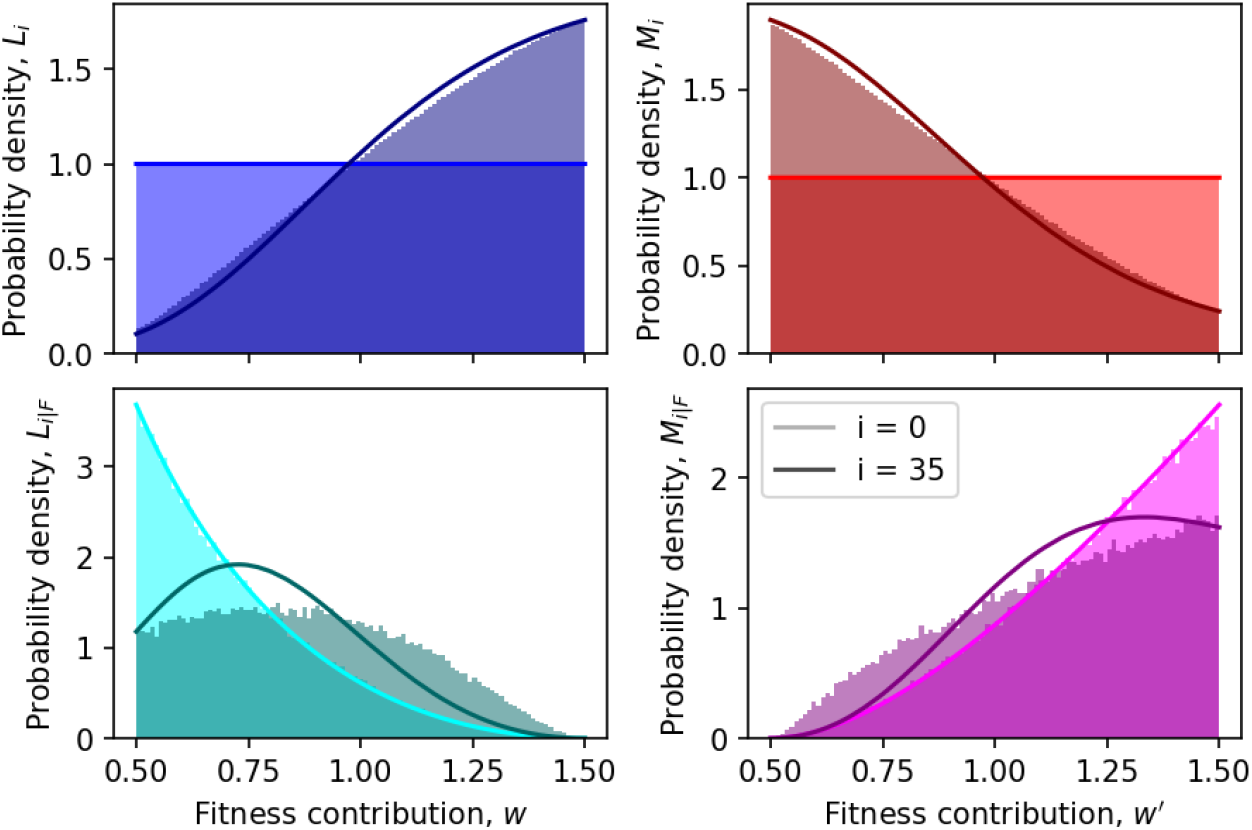
Simulation results (colored histograms) and analytical approximations (solid lines) for the density functions *L*_*i*_ (blue), *L*_*i*|*F*_ (cyan), *M*_*i*_ (red) and *M*_*i*|*F*_ (magenta) in a multiplicative fitness landscape. The initial distributions (at *i* = 0) are in lighter colors, while the distributions for *i* = 35 are in darker colors. Results for *B* = 2, *N* = 100, *L*_0_ = *M*_0_ = *U* (0.5, 1.5) are shown.

**Figure S.5:**
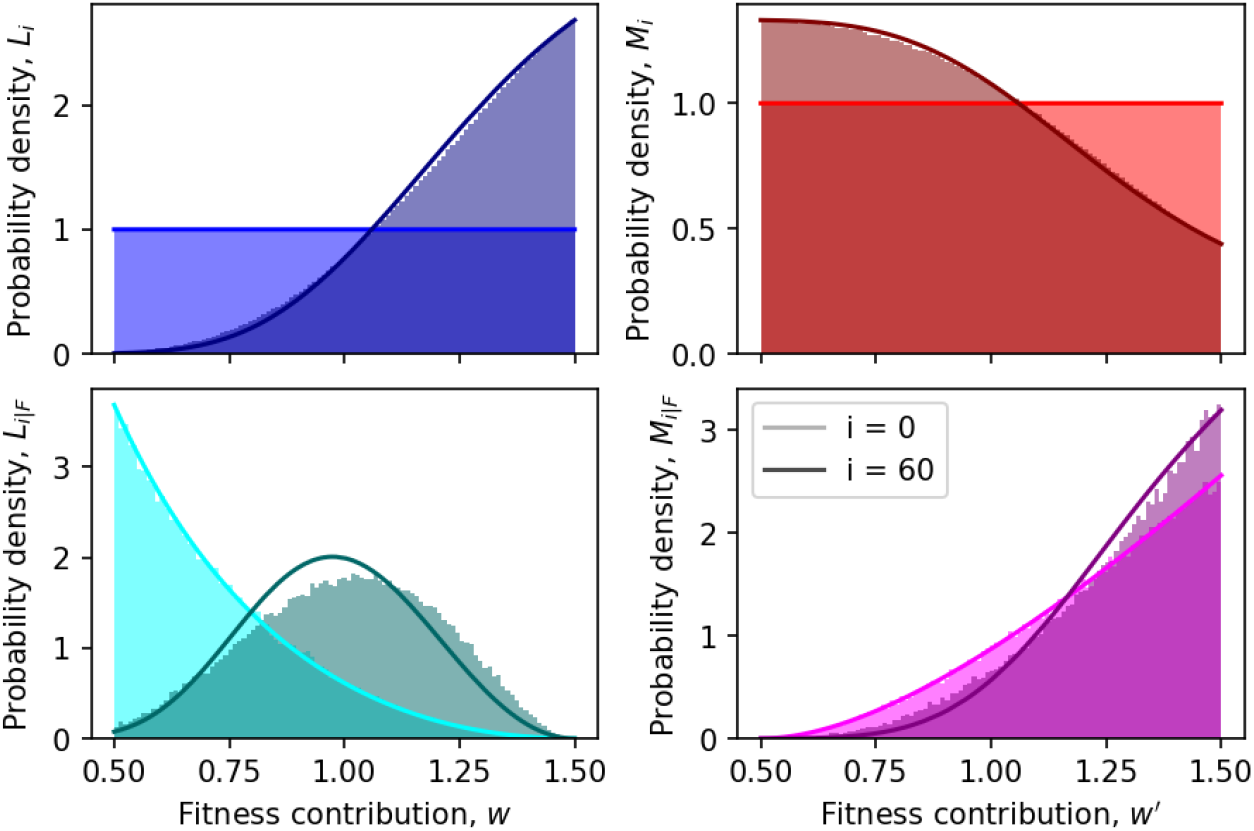
Simulation results (colored histograms) and analytical approximations (solid lines) for the density functions *L*_*i*_ (blue), *L*_*i*|*F*_ (cyan), *M*_*i*_ (red) and *M*_*i*|*F*_ (magenta) in a multiplicative fitness landscape. The initial distributions (at *i* = 0) are in lighter colors, while the distributions for *i* = 60 are in darker colors. Results for *B* = 4, *N* = 100, *L*_0_ = *M*_0_ = *U* (0.5, 1.5) are shown.

**Figure S.6:**
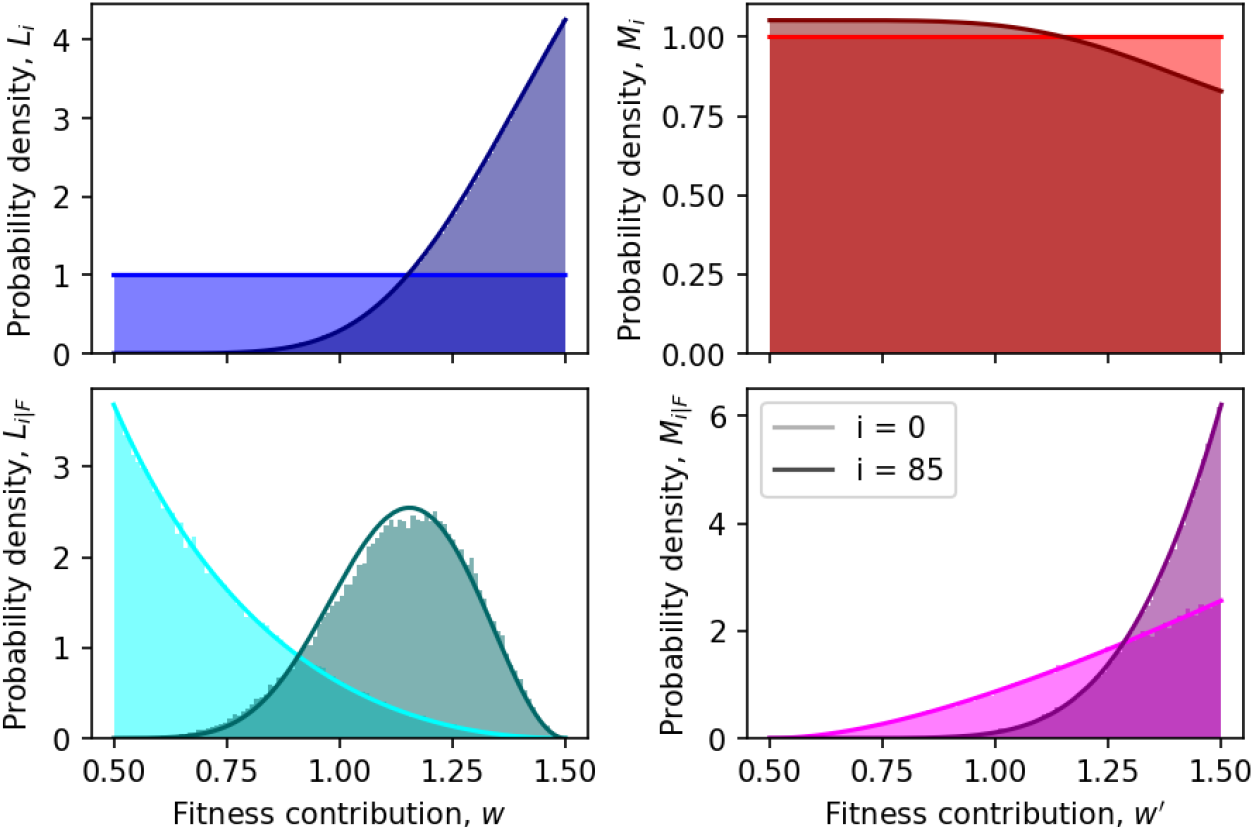
Simulation results (colored histograms) and analytical approximations (solid lines) for the density functions *L*_*i*_ (blue), *L*_*i*|*F*_ (cyan), *M*_*i*_ (red) and *M*_*i*|*F*_ (magenta) in a multiplicative fitness landscape. The initial distributions (at *i* = 0) are in lighter colors, while the distributions for *i* = 85 are in darker colors. Results for *B* = 20, *N* = 100, *L*_0_ = *M*_0_ = *U* (0.5, 1.5) are shown.

**Figure S.7:**
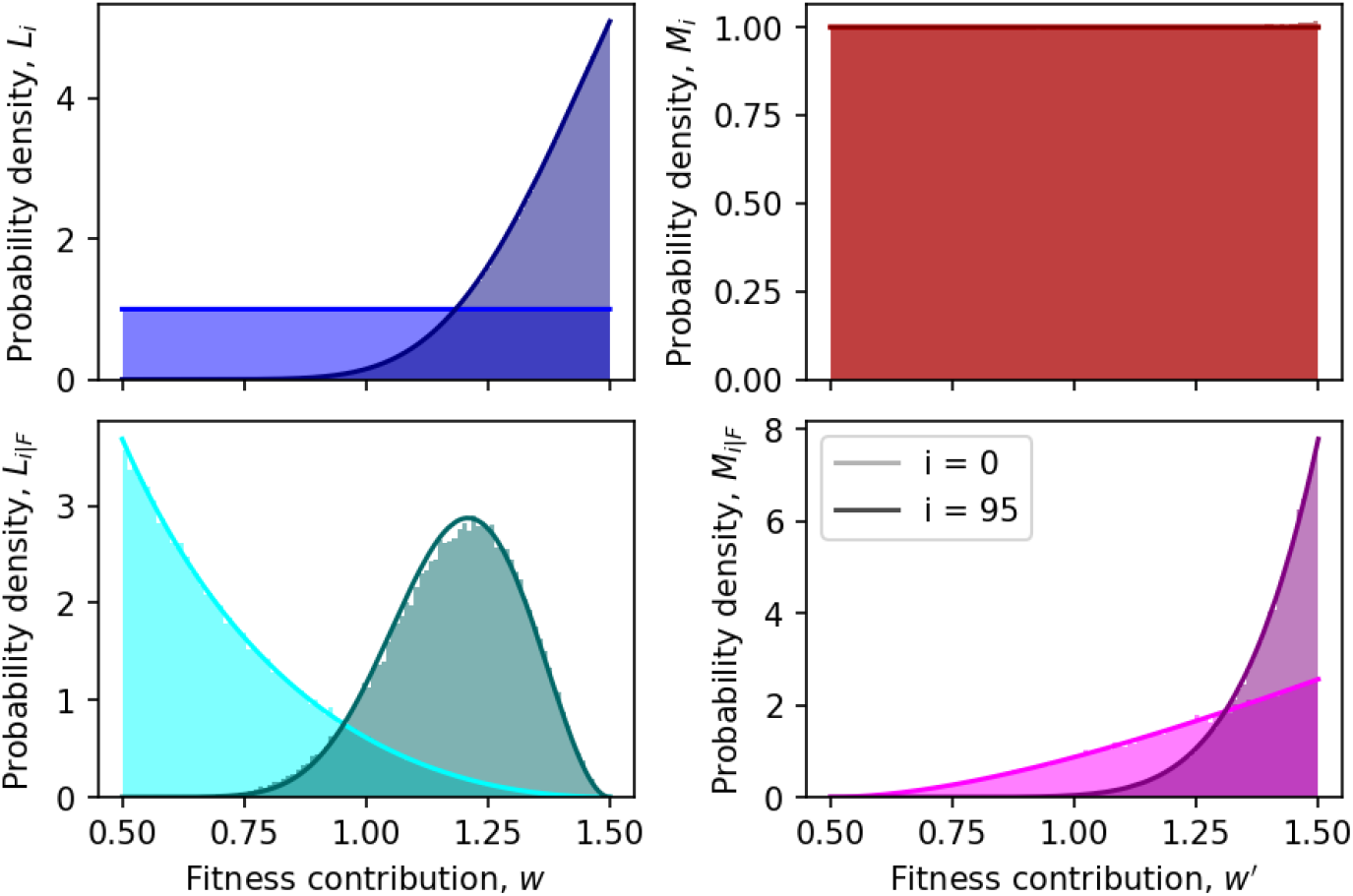
Simulation results (colored histograms) and analytical approximations (solid lines) for the density functions *L*_*i*_ (blue), *L*_*i*|*F*_ (cyan), *M*_*i*_ (red) and *M*_*i*|*F*_ (magenta) in a multiplicative fitness landscape. The initial distributions (at *i* = 0) are in lighter colors, while the distributions for *i* = 95 are in darker colors. Results for an infinite alleles model, *N* = 100, *L*_0_ = *M*_0_ = *U* (0.5, 1.5) are shown.

**Figure S.8:**
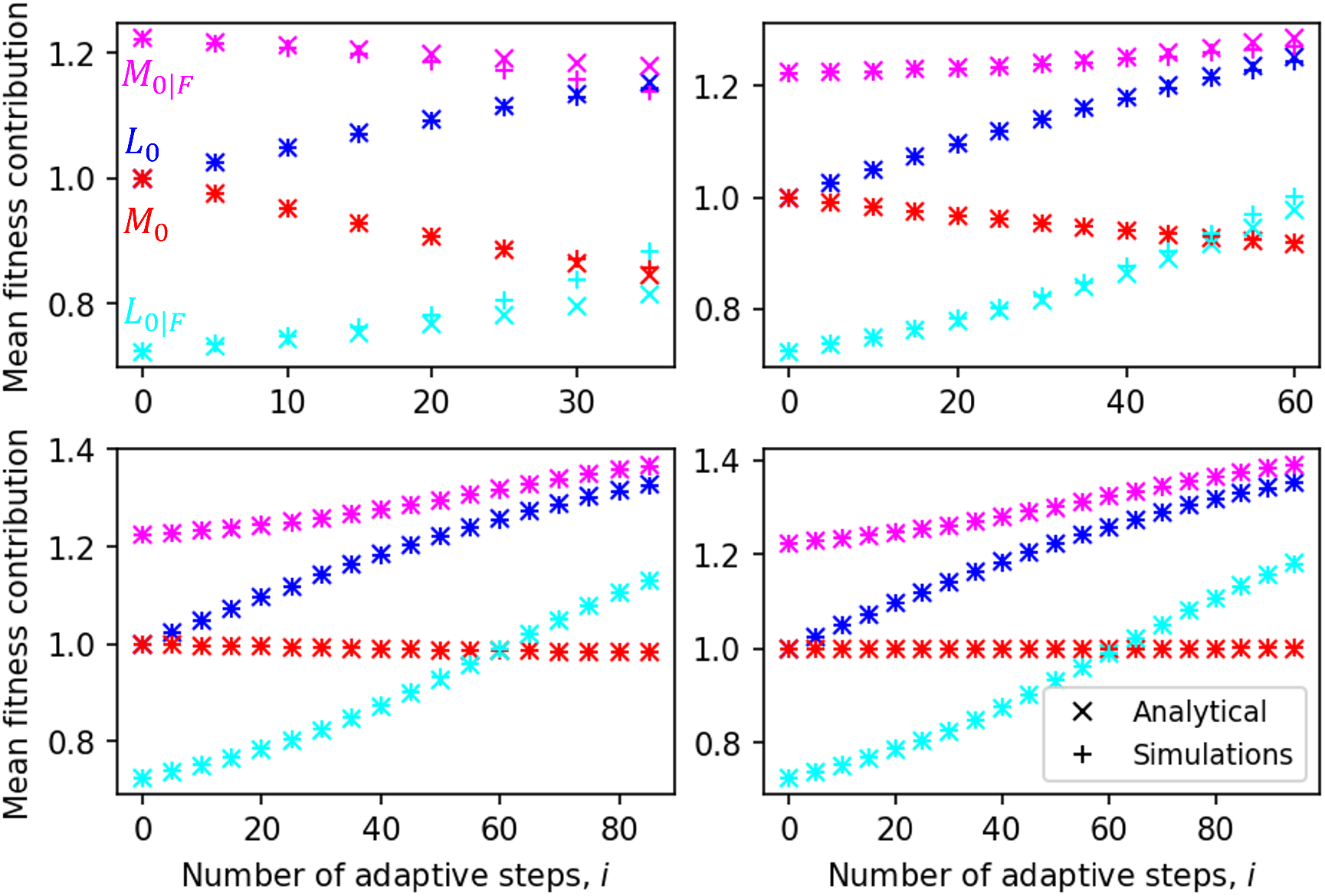
Analytical approximations (crosses) and simulation results (plus signs) for a multiplicative fitness landscape for the four distributions for the fitness contributions: *L*_*i*_ (blue), *L*_*i*|*F*_ (cyan), *M*_*i*_ (red) and *M*_*i*|*F*_ (magenta), for different number of available alleles at each site, *B*. In each case, *i* steps in the adaptive walk are taken such that the beneficial fraction of the DFE falls below 15%. For *B*=2 (top left), 4 (top right), 20 (bottom left) and the infinite alleles model (bottom right), the number of steps taken are *i*=35, 60, 85 and 95, respectively. Results are shown for *N* = 100 and *L*_0_ = *M*_0_ = *U* (0, 1).

**Figure S.9:**
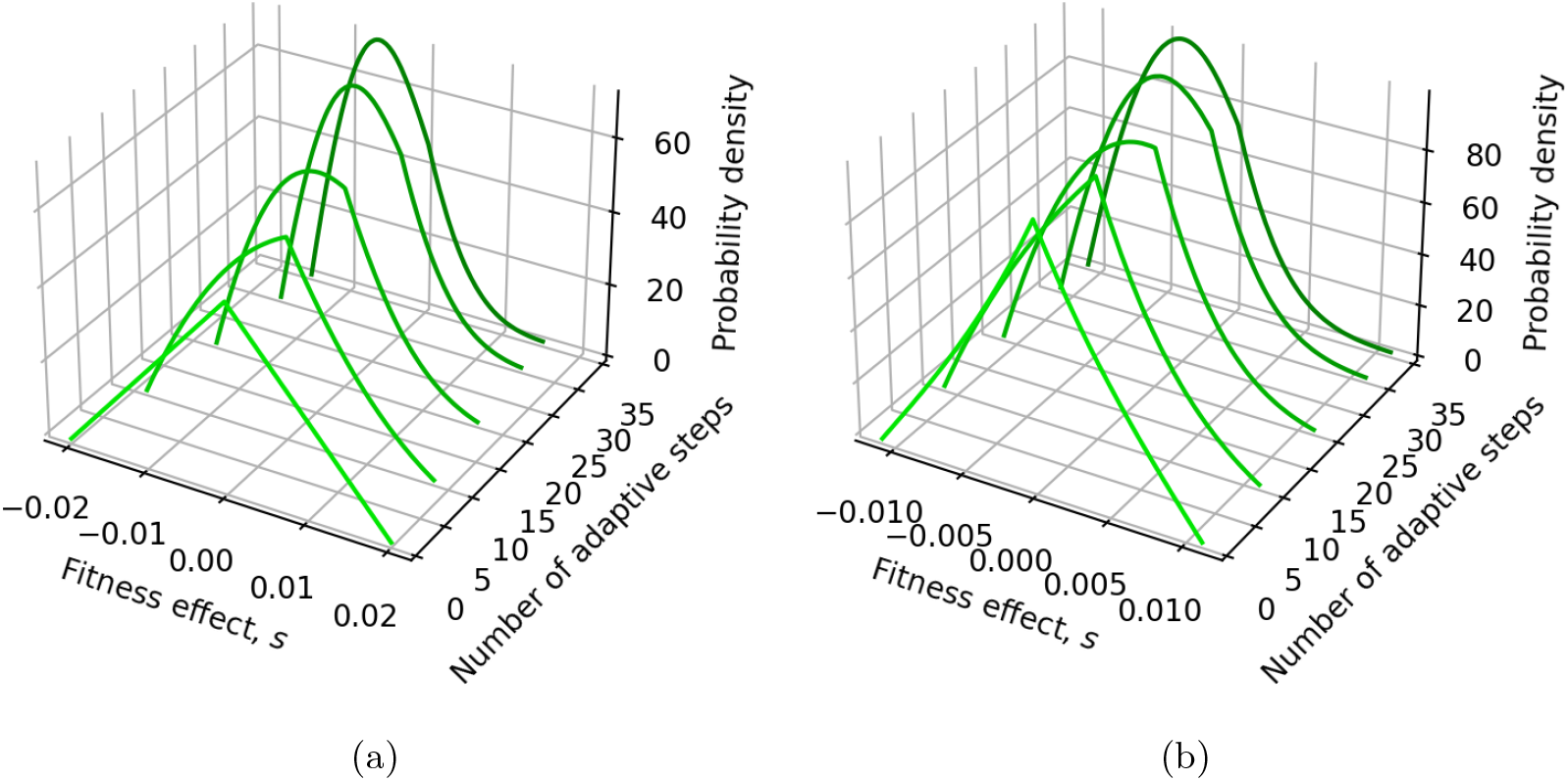
The DFE changes shape as the number of steps in the adaptive walk increases. (a) Analytical results for an additive landscape and initial fc distributions *L*_0_ = *M*_0_ = *U* (0, 1). (b) Analytical results for a multiplicative landscape and initial fc distributions *L*_0_ = *M*_0_ = *U* (0.5, 1.5). Results are shown for *N* = 100 and *B* = 2.

**Figure S.10:**
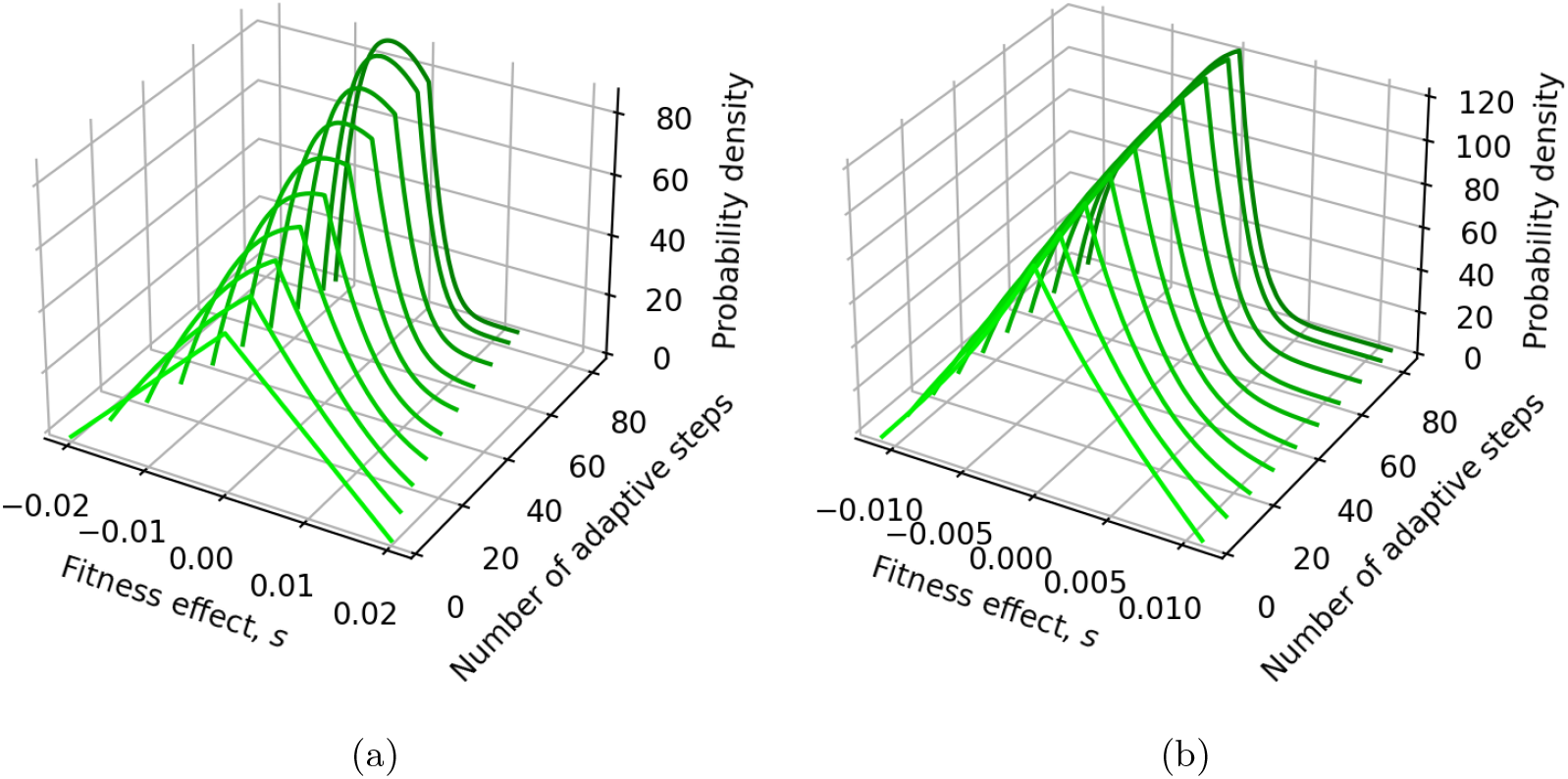
The DFE changes shape as the number of steps in the adaptive walk increases. (a) Analytical results for an additive landscape and initial fc distributions *L*_0_ = *M*_0_ = *U* (0, 1). (b) Analytical results for a multiplicative landscape and initial fc distributions *L*_0_ = *M*_0_ = *U* (0.5, 1.5). Results are shown for *N* = 100 and *B* = 20.

**Figure S.11:**
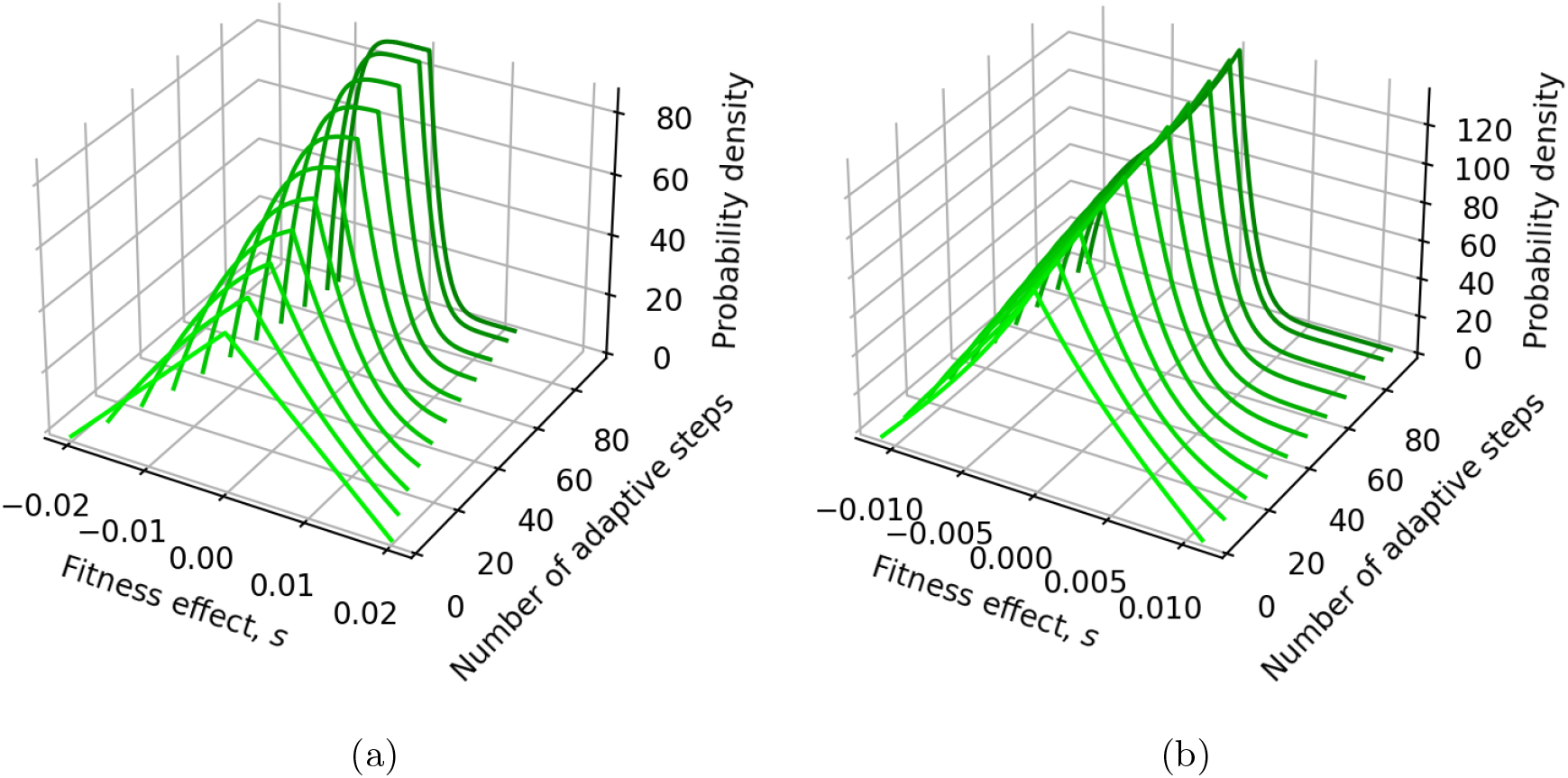
The DFE changes shape as the number of steps in the adaptive walk increases. (a) Analytical results for an additive landscape and initial fc distributions *L*_0_ = *M*_0_ = *U* (0, 1). (b) Analytical results for a multiplicative landscape and initial fc distributions *L*_0_ = *M*_0_ = *U* (0.5, 1.5). Results are shown for *N* = 100 and an infinite alleles model.

### S.2 Dynamics of a highly adapted population

To study a well-adapted population, we analyzed the case when the fcs of the alleles that comprise the initial genome tend to be larger than the fcs of available mutations, for the case of an additive fitness landscape. This was performed by changing the initial distributions *L*_0_ and *M*_0_ from uniform distributions to distributions that mimic an adapted population. In this case, we used simple linear density functions on (0,1): *L*_0_(*w*) = 2*w* and *M*_0_(*w*^′^) = 2 − 2*w*^′^, such that the focal alleles have a higher fitness on average while available mutations tend to be less fit, as shown in Figure S.12. Adaptive walks were performed until the beneficial fraction of the DFE fell bellow 50% of its initial value (that is, from a beneficial fraction of 1*/*6 to beneficial fractions lower than 1*/*12), which corresponds to 10, 17, 30 and 35 mutations for *B* = 2, 4, 20 and an infinite number of alleles, respectively.

Figure S.13 shows the analytical results for the DFE as adaptation proceeds, for an initially well adapted population. We note that the resulting DFEs have a more realistic shape (compare Figure 5 and Figure S.14, with Cotto and Day (2023), and Kassen and Bataillon (2006)). As before, the analytical approximation agrees well with simulation results for a large number of alleles, and gradually loses accuracy if *B* is decreased (Figure S.14). Interestingly, for this population we observe similar results as described previously: the dynamics for *L*_0_ and *M*_0_ are symmetrical for *B* = 2, while for higher values of *B* the distributions of available mutations remain almost unchanged throughout the walk.

**Figure S.12:**
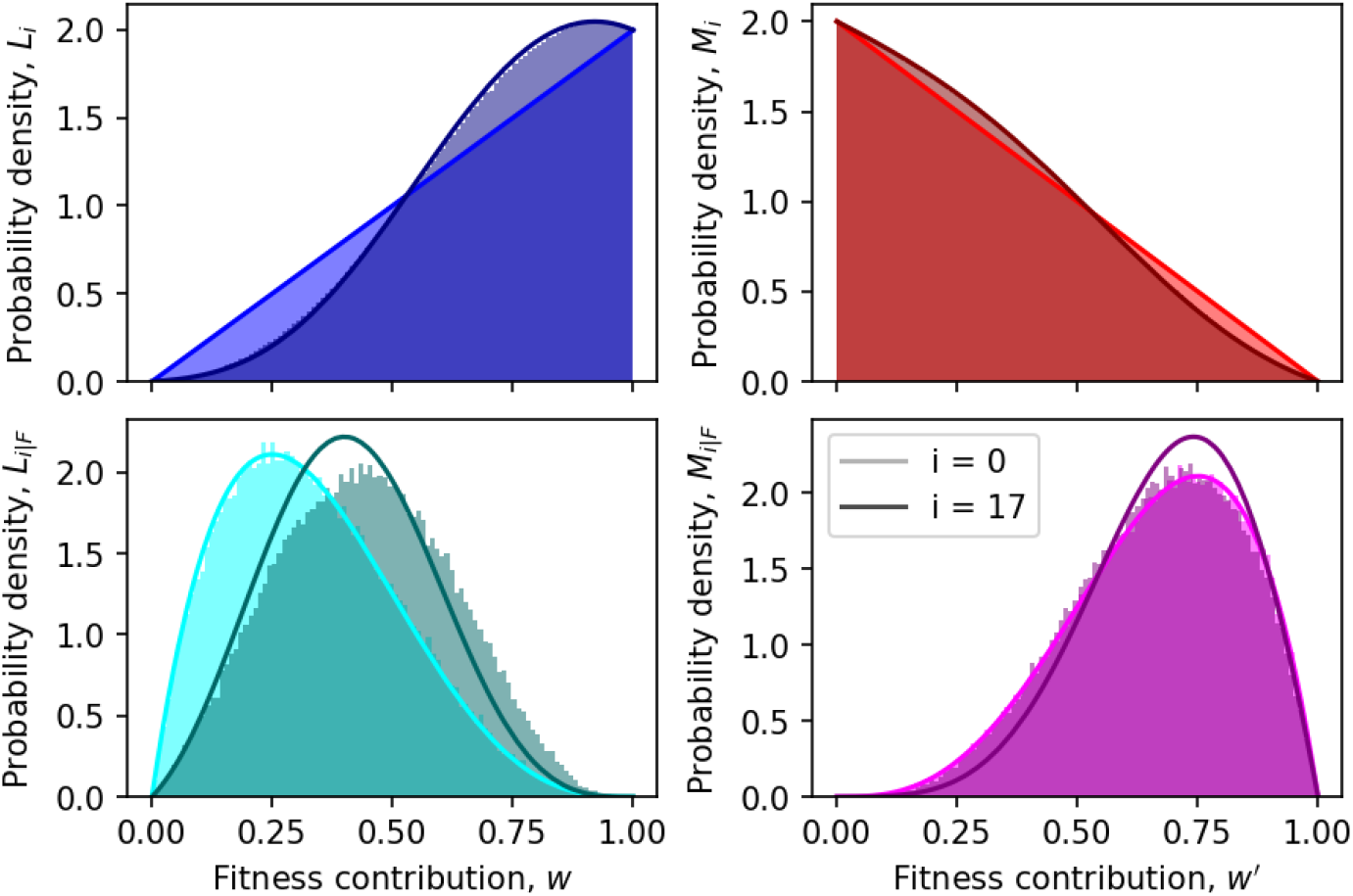
Simulation results (colored histograms) and analytical approximations (solid lines) for the density functions *L*_*i*_ (blue), *L*_*i*|*F*_ (cyan), *M*_*i*_ (red) and *M*_*i*|*F*_ (magenta) in a additive fitness landscape. The initial distributions (at *i* = 0) are in lighter colors, while the distributions for *i* = 17 are in darker colors. Results for *B* = 4, *N* = 100, *L*_0_(*w*) = 2*w* and *M*_0_(*w*^′^) = 2 − 2*w*^′^ are shown.

**Figure S.13:**
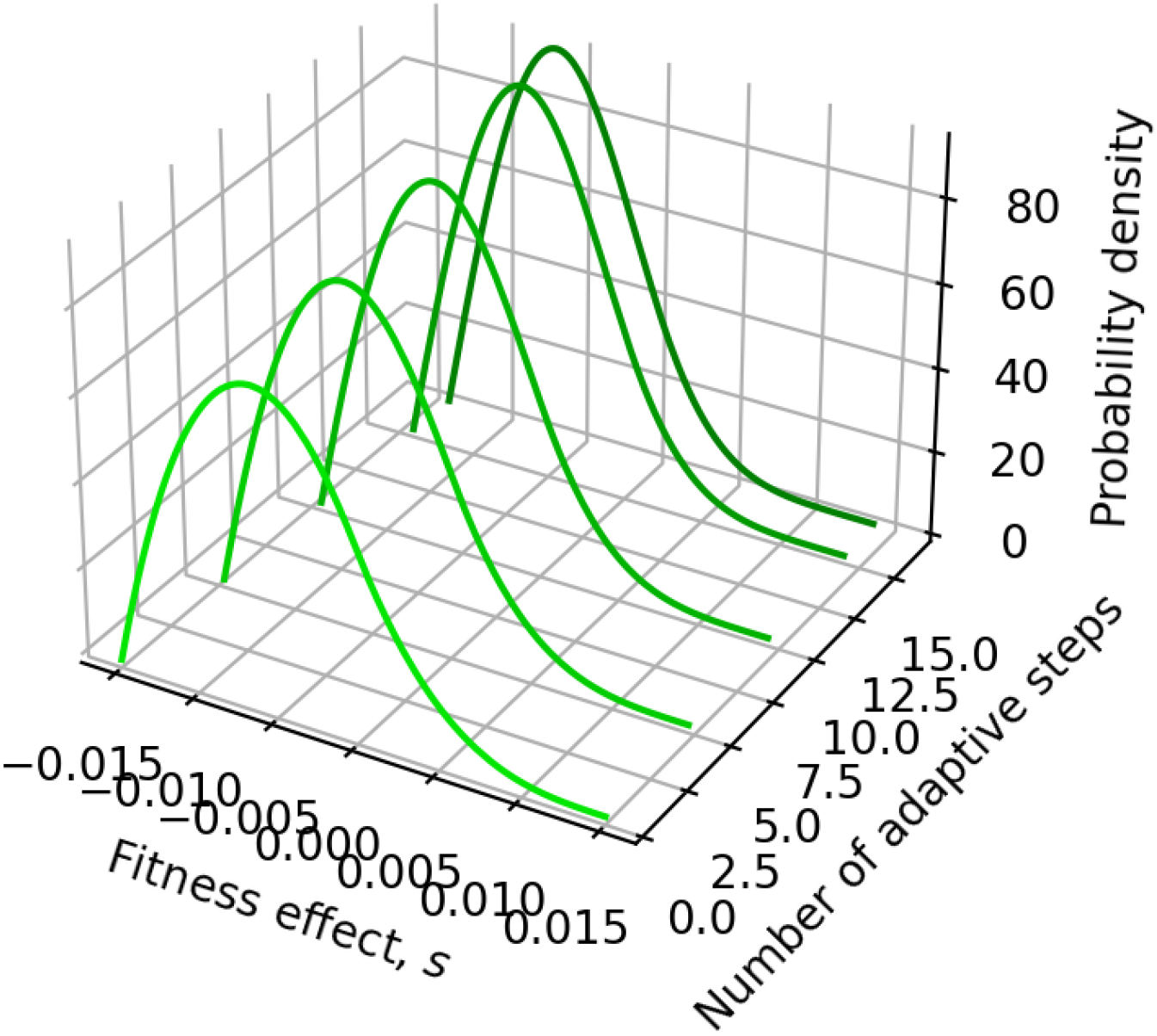
The DFE changes shape as the number of steps in the adaptive walk increases. Analytical results for an additive landscape and initial distributions *L*_0_(*w*) = 2*w* and *M*_0_(*w*^′^) = 2 − 2*w*^′^. Results shown for *N* = 100 and *B* = 4.

**Figure S.14:**
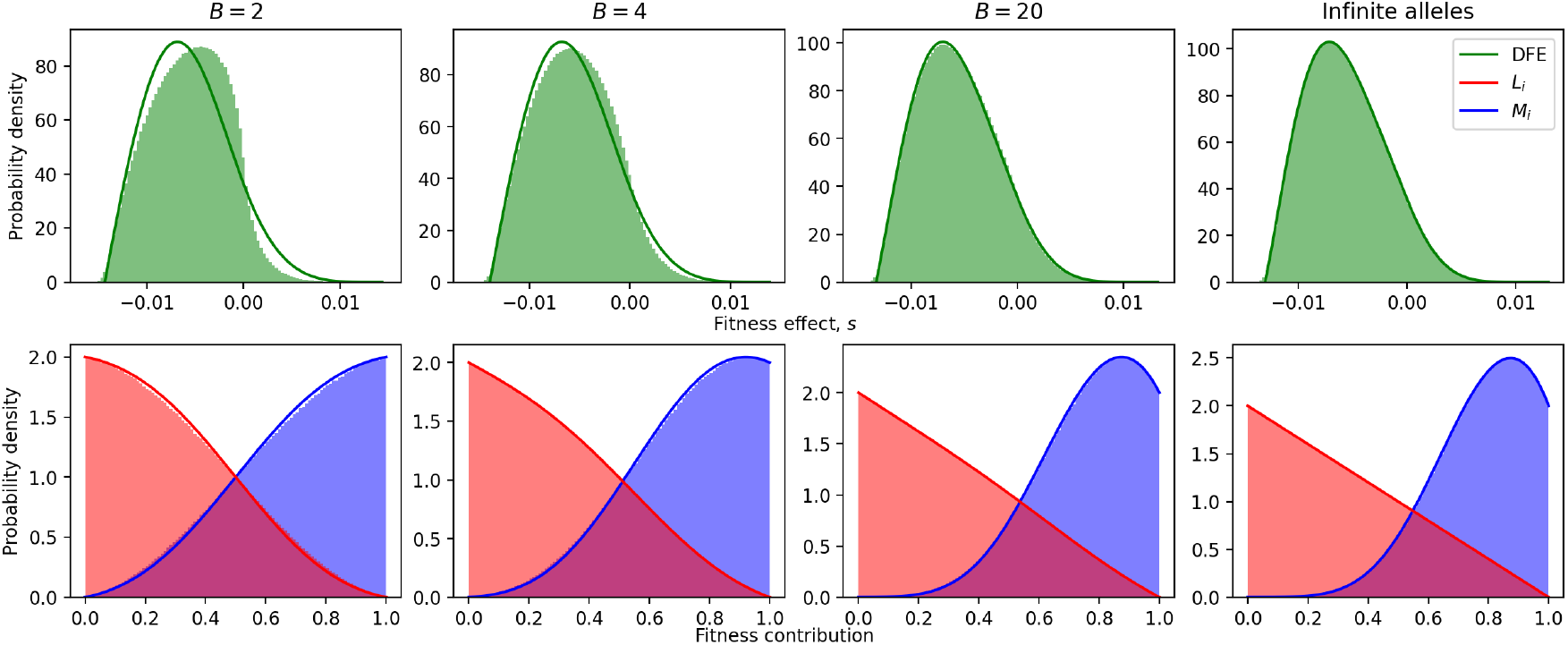
Analytical approximations (solid lines) and simulation results (histograms) for the DFE (top row) and distributions *L*_*i*_ and *M*_*i*_ (bottom row), for different number of available alleles at each site, *B*, for an additive fitness landscape. In each case, the number of steps in the adaptive walk are such that the beneficial fraction of the DFE falls just below half the initial beneficial fraction. For *B*=2, 4, 20 and the infinite alleles model, the numbers of steps taken are 10, 17, 30 and 35, respectively. Despite the near-constant beneficial fraction of the DFE, the distributions of available fitness contributions differ significantly across cases. Results are shown for *N* = 100, *L*_0_(*w*) = 2*w* and *M*_0_(*w*^′^) = 2 − 2*w*^′^.

### S.3 Effects of Assumptions 1 and 2

Here, we address the discrepancies in the DFE predicted by our analytical approximations (see Figures 5 and 6). To showcase these observations in further detail, results for the analytical and simulated DFEs are shown in Figure S.15, for *B* = 4 and up to 50 adaptive steps. We can see two main discrepancies: the truncation of the elongated tails of the distribution (see inset); and the underestimation of deleterious mutations with small fitness effect.

First, we consider the elongated tails in the simulation results, which are truncated in the analytical approximation. This discrepancy arises from Assumption 1: that the fc of any site is independent of the overall fitness (which is approximated as a constant). The distribution tails arise from simulation samples in which the fitness is lower than average, increasing the absolute value of the fitness effect (see Equation 4).

To demonstrate this effect, we derived the analytical prediction for the initial DFE without imposing Assumption 1. In this case, we take fitness, *W*, to be a random variable. Considering a mutation that changes the fc of the *k*th site from *w* to *w*^′^, let 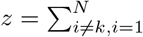 be the random variable for the sum of the fc of the *N* − 1 unmutated sites, such that the fitness is given by

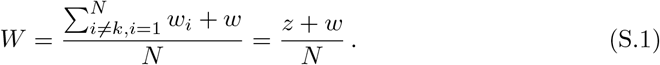

The fitness effect of the mutation can be written from Equation 4, in terms of *w, w*^′^ and *z*,

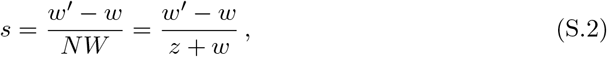

and then the DFE can be computed as the convolution of these three independent random variables. Following Equation 13, if we know the pdf for the random variable *z* (here denoted *f* (*z*)), the DFE is given by:

**Figure S.15:**
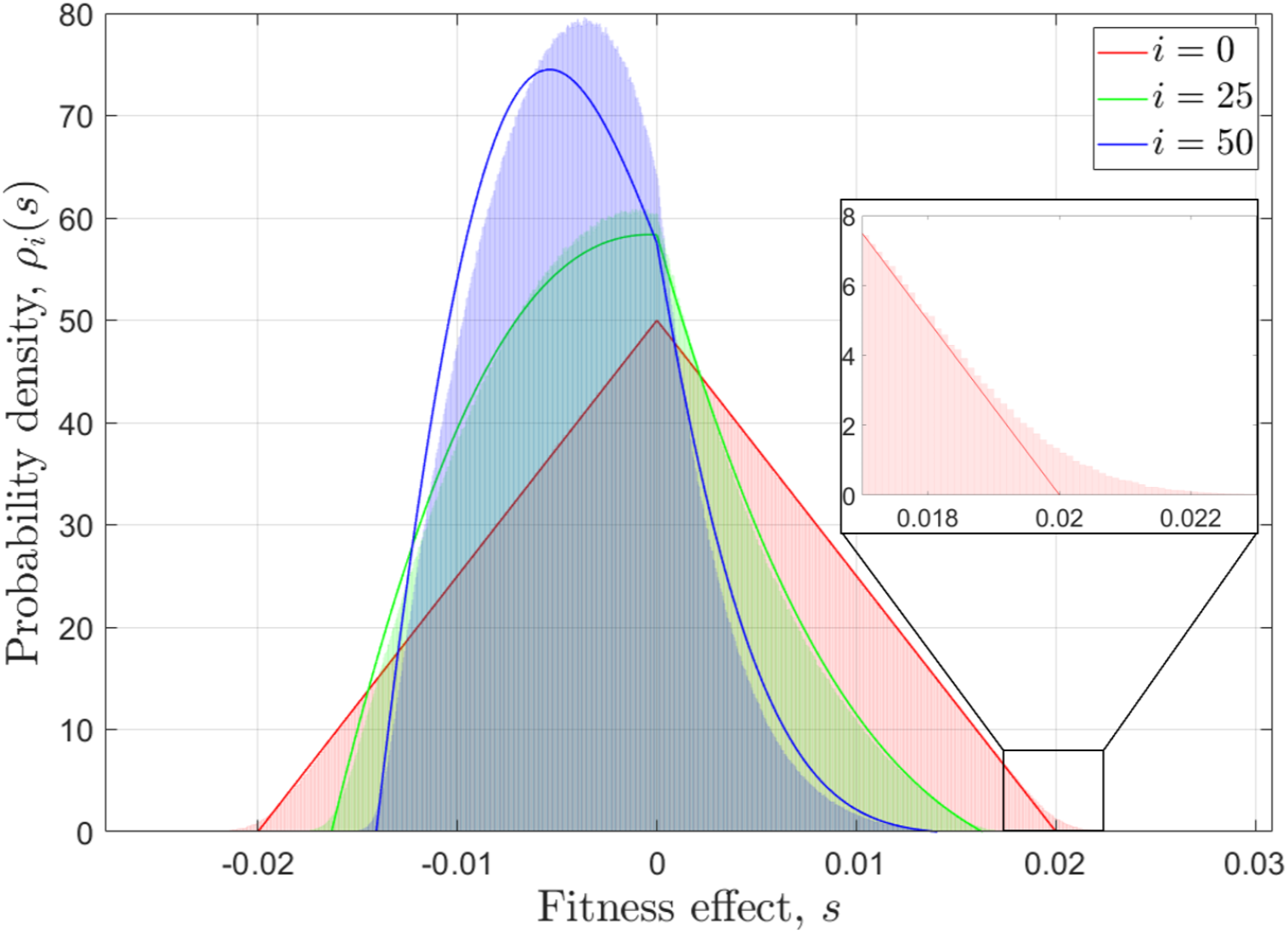
DFEs for different number of fixed mutations, *i*. Here analytical approximations are shown with solid lines and simulation results appear as colored histograms. Results for *N* = 100, *B* = 4, *L*_0_ = *M*_0_ = *U* (0, 1) and 10^5^ adaptive walks are shown.

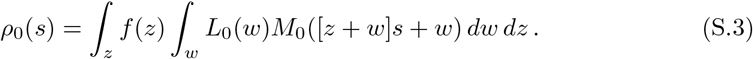

Although computing *f* (*z*) is generally complicated, this is not the case for our initial uniform distributions, *L*_0_ = *M*_0_ = *U* (0, 1). In this case *f* (*z*) is described by the Irwin-Hall distribution, defined as the pdf of the sum of a number of uniformly distributed random variables (here, the sum of *N* − 1 such variables). Applying this to Equation S.3, we arrive at the solid line shown in Figure S.16. Thus the full analytical prediction, in which Assumption 1 is relaxed, matches the tails of the simulation results; this holds even when we consider only a modest number of sites, *N* = 10, for which the tails of the distribution are longer.

Unfortunately, extending this approach to further steps in the walk or to more complicated initial distributions proves unwieldy, thus Assumption 1 was imposed to provide a reasonable simplification.

We now consider the other main discrepancy between simulations and predictions from Figures 5 and 6. In particular, the analytical results underestimate the density of deleterious mutations with small |*s*| value. This is largely due to the fact that the random variables *w* and *w*^′^ are not independent once a mutation fixes, which breaks Assumption 2. As described in the main text, this dependency emerges at sites where a previous fixation has occurred, such that the fc of the site is higher than average (since a mutation fixed there), while the fc of one available mutation (the one that reverses the fixation) is lower than average. This produces a dependence in which higher fcs at the site, *w*, are correlated with lower fcs in the available mutations, *w*^′^.

It is difficult to demonstrate, analytically, the effect of this assumption on the predicted DFE, since computing a DFE which retains the dependencies between *w* and *w*^′^ at each step *i* is somewhat complex. Instead, we can manipulate the simulation algorithm to manually impose the assumption of independence. In other words, we were able to simulate “fake” DFEs in which Assumption 2 holds.

**Figure S.16:**
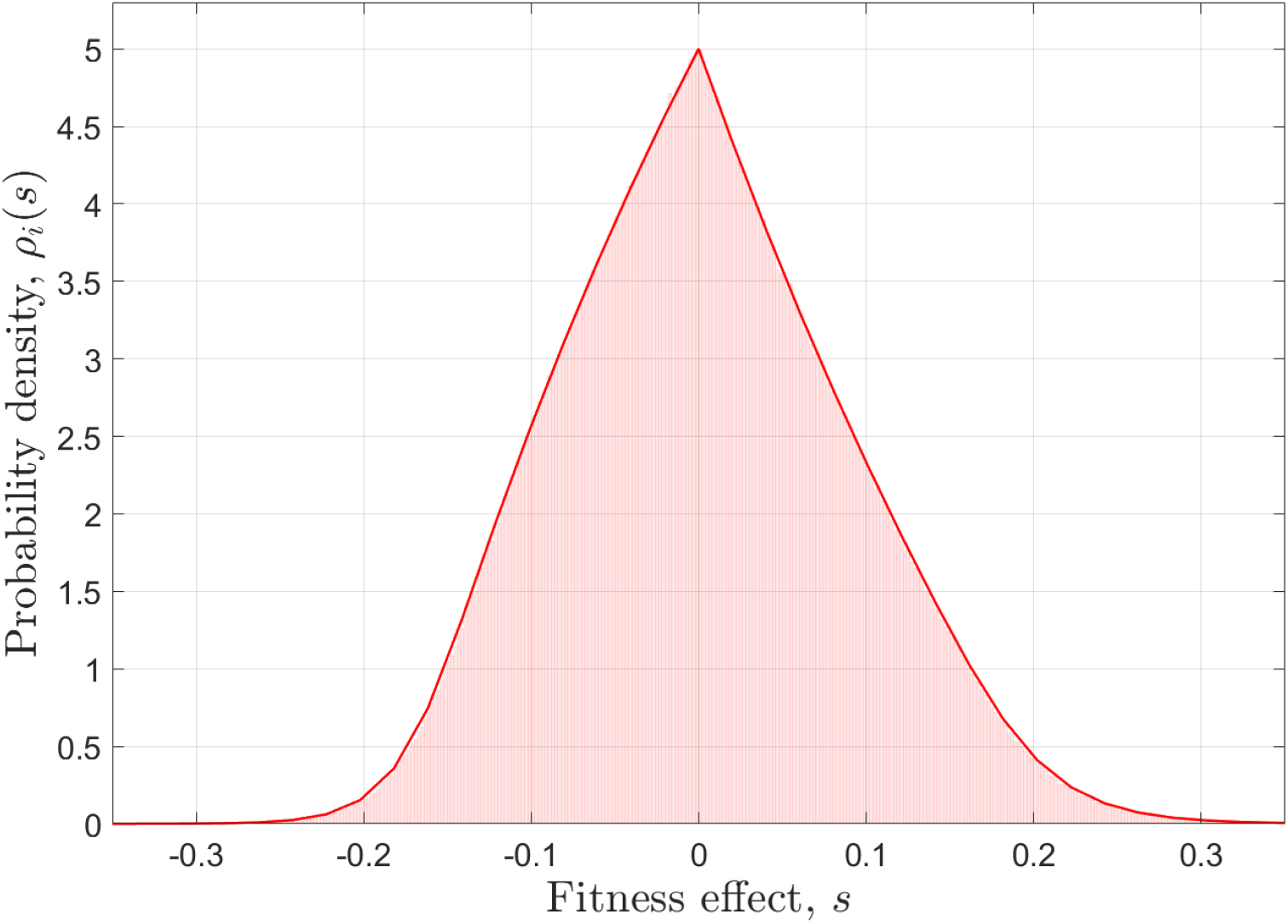
DFE at the beginning of the walk, *ρ*_0_(*s*). The corrected analytical approximation is shown as a solid line and simulation results as a colored histogram. Results for *N* = 10, *L*_0_ = *M*_0_ = *U* (0, 1) and 10^5^ adaptive walks are shown.

In all the DFEs in the main text, we consider each of *N* sites in turn, then consider each of the *A* possible mutations at that site, and compute a single fitness effect, *s*, for each of these mutations. To create a DFE in which Assumption 2 holds, we instead choose a site at random, and then choose an available mutation <u>from any site in the genome at random</u>, until we have the same total of *NA* fitness effects in the DFE. In this way both the fcs *w* and *w*^′^ are random, uncorrelated samples, regardless of the sites at which they might occur.

In Figure S.17 we demonstrate the much improved agreement between our previous analytical predictions and the DFEs thus obtained when Assumption 2 is imposed. As expected, once the dependency of *w* and *w*^′^ is removed (by sampling the fc from different sites) the overall shape of the simulated DFE closely matches the analytical prediction. In this case the analytical prediction still truncates the tails of the distribution, and thus predicts slightly higher densities for the bulk of the distribution.

**Figure S.17:**
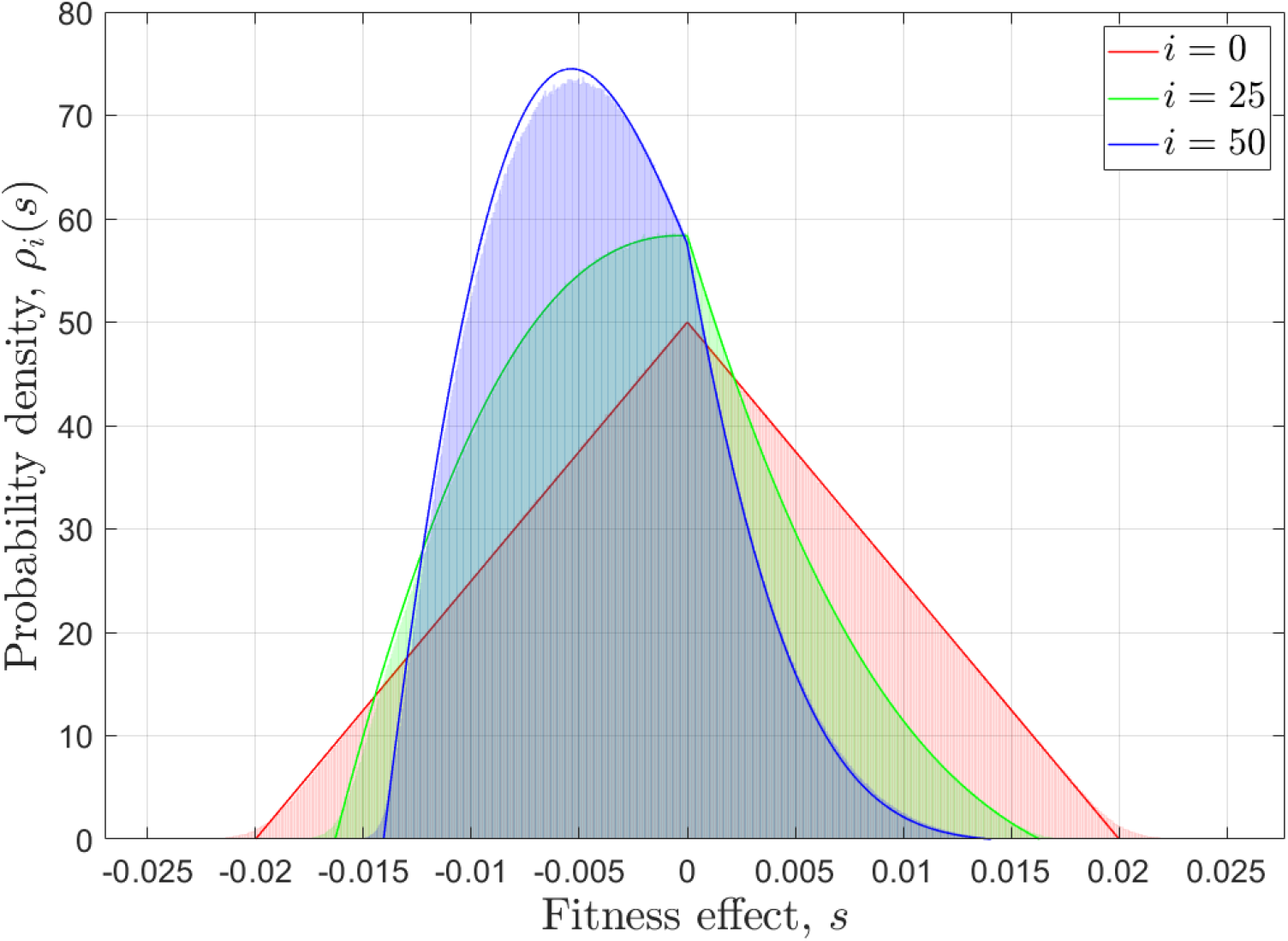
DFEs for different number of fixed mutations, *i*, when Assumption 2 is imposed. Here analytical approximations are shown with solid lines and simulation results appear as colored histograms. Results for *N* = 100, *B* = 4, *L*_0_ = *M*_0_ = *U* (0, 1) and 10^5^ adaptive walks are shown.

### S.4 Fitness over time

A common observation in any adaptive process is the diminishing returns on the increment of fitness over time, observed experimentally by Lenski (2017) for long periods of time. We studied this observation for our adaptive walks, for analytical approximations and simulation results, as shown in Figure S.18, where the mean fitness at each adaptive step is plotted against the average number of spontaneous mutations taken to achieve that step (in log space). As before, we considered additive and multiplicative fitness landscapes, and results are very similar for both models. We observe the characteristic diminishing returns in the increment of fitness, due to the depletion of mutations with high fitness effect from the genetic pool. Although the analytical approximations for the high-*B* replicates are very close to simulation results, for *B* = 4 and *B* = 2 the analytical predictions grossly underestimate the times between fixations. The discrepancies arise from a lower availability of beneficial mutations in the simulations with respect to the analytical approximations: in simulated walks, each site can only access *B* possible alleles, thus if a site has had a successful mutation, it will be unlikely for it to have other advantageous alleles to chose from. In contrast, in the analytical approximations each site has the same chance of mutating to a fitter allele, generating faster fixations if random mutations occur in large quantities.

**Figure S.18:**
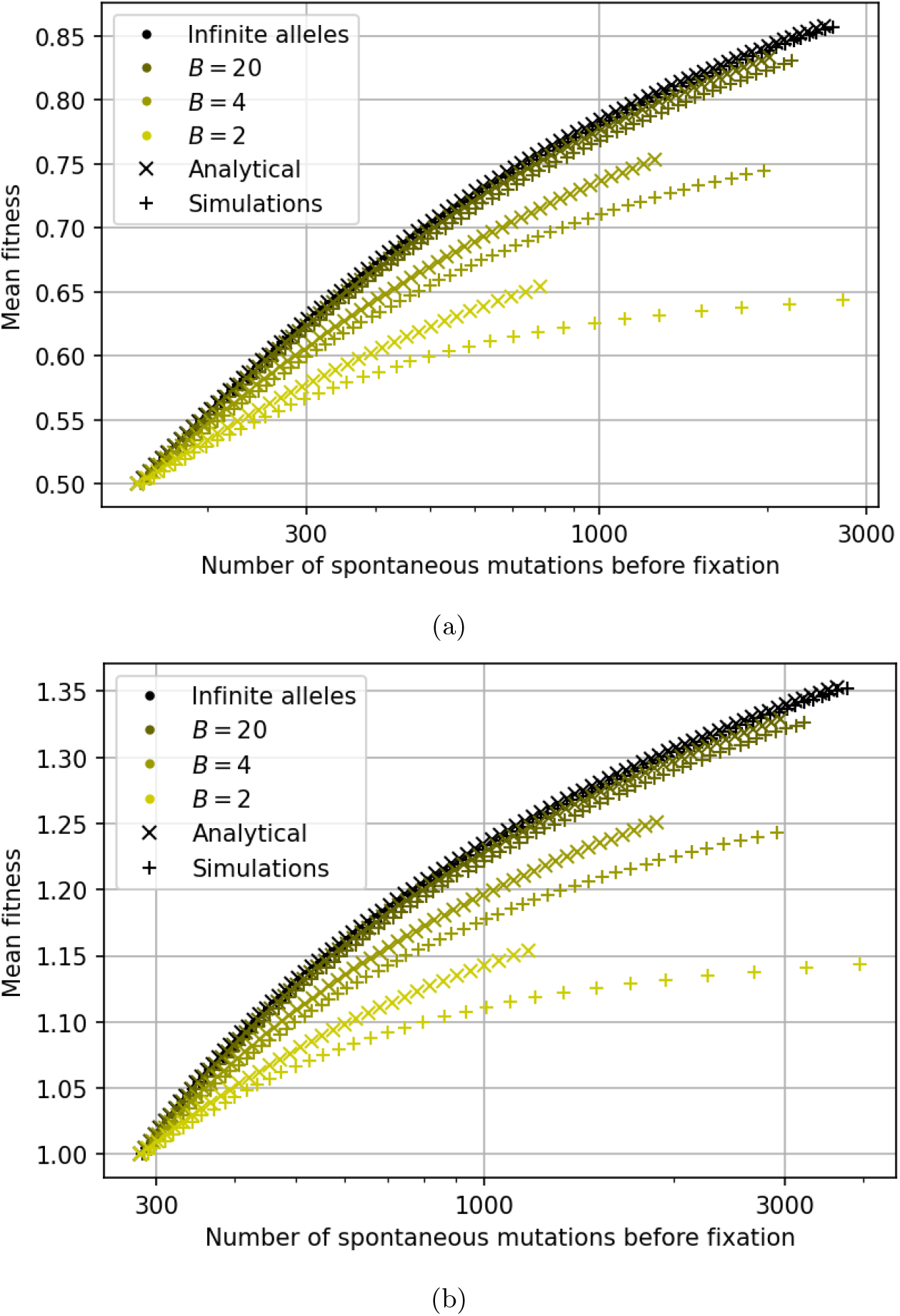
Analytical approximations (crosses) and simulation results (plus signs) for the mean fitness at each adaptive step versus the number of spontaneous mutations that have occurred (whether or not they fixed), for different number of available alleles at each site, *B*. In each case, a symbol represents a step and a total of *i* steps in the adaptive walk are taken such that the beneficial fraction of the DFE falls below 15%. For *B*=2, 4, 20 and infinite model, the number of steps taken are *i* = 35, 60, 85 and 95, respectively. Note that we have a log horizontal axis. Results are shown for *N* = 100 and (a) an additive fitness landscape (*L*_0_ = *M*_0_ = *U* (0, 1)) and (b) a multiplicative fitness landscape (*L*_0_ = *M*_0_ = *U* (0.5, 1.5)).

